# Plant-associated *Streptomyces* detoxify the mycotoxin fusaric acid by amino acid conjugation

**DOI:** 10.64898/2026.08.26.747244

**Authors:** Emtinan Diab, Chao Du, Simon C. Verdel, Mees R. Kunnen, Rachel Stuij, Somayah S. Elsayed, Jos M. Raaijmakers, Gilles P. van Wezel

## Abstract

Streptomycetes are prevalent members of soil and plant microbiomes, yet how they cope with toxins produced by root-infecting fungal pathogens remains poorly understood. Plant pathogenic *Fusarium* species produce the mycotoxin fusaric acid (FA) that contributes to virulence and perturbs rhizosphere microbiome dynamics. Here, we show that root-colonising *Streptomyces* sp. ATMOS43 neutralizes FA through amino acid conjugation. Metabolomics revealed the formation of single amino acid and dipeptidyl conjugates of FA, with FA-Ser as a major conjugate that lacked detectable toxicity in *in vitro* and *in planta* assays. Proteomics and physiological analyses revealed that FA toxicity involves, in part, zinc chelation, which is abolished upon conjugation of FA to Ser. Co-cultivation experiments further showed that *Streptomyces* sp. ATMOS43 restores growth of FA-sensitive streptomycetes, indicating that conjugation can mitigate the impact of FA on plant microbiome assembly. Together, our findings show that plant-associated streptomycetes can protect plants by directly inhibiting *Fusarium* growth and by neutralizing its toxic virulence factor FA.

## INTRODUCTION

The growing global population puts increased pressure on crop productivity and food security. Major crops are increasingly threatened by plant pathogens due to climate change, prompting the need for effective and sustainable alternatives. Plants live in close association with taxonomically diverse microbiota that shape plant physiology, growth and health [1]. In return, plants provide nutritionally rich habitats for microbial colonization, both in the rhizosphere and endophytic compartments. Microbes colonizing plant tissues exploit plant-derived nutrients for growth and to establish commensal, mutualistic or pathogenic interactions depending on the host plant species and environmental context [2]. These interactions are strongly shaped by specialized metabolites produced by both partners: plant metabolites contribute to defence and microbiome assembly by selectively promoting or inhibiting microbial taxa, while microbial metabolites support colonization, competition, or virulence [3–5]. Hence, the rhizosphere and endosphere have long been recognized as a microbial ’playground’ and ’battlefield’ [6].

Actinobacteria are prevalent members of the rhizosphere and endosphere microbiomes of many plant species [7, 8]. Streptomycetaceae—a family within the phylum Actinobacteria—comprises multicellular mycelial bacteria that undergo a complex mycelial lifecycle and reproduce through sporulation [9, 10]. These bacteria are known as Nature’s medicine makers, with a huge capacity to produce antibiotics, antifungals and other bioactive natural products [11–13]. Streptomycetes are well known for their ability to promote plant growth and suppress plant diseases [14–16]. For example, several *Streptomyces* strains have been reported to suppress *Fusarium* infections, typically through the production of antifungal natural products [15, 17, 18].

*Fusarium* species are among the major fungal threats to crop production worldwide, causing diseases such as root rot, head blight and vascular wilt [19–21]. Current control measures of *Fusarium* infections are hampered by the rapid emergence of fungicide resistance [22]. Hence there is an urgent need for novel measures to control *Fusarium*. In this work, we investigated whether plant-associated streptomycetes can mitigate *Fusarium* pathogenicity through neutralization of the mycotoxin fusaric acid (FA). Pathogenicity of *Fusarium* is associated with the production of FA and other mycotoxins such as trichothecenes and zearalenone [21, 23]. FA (5-butylpicolinic acid) is produced by multiple *Fusarium* species and its production is associated with virulence [24]. In plants, FA induces oxidative stress, disrupts mitochondrial function, and damages proteins and DNA, ultimately leading to cell death [25–27]. FA also affects the composition of the rhizosphere microbial community by inducing changes in root exudation [28]. Moreover, differential susceptibility to FA among plant-associated microorganisms can shape the ecology of microbe–microbe interactions in the rhizosphere [29]. To date, however, the fundamental mechanisms underlying FA susceptibility and tolerance of plant-associated microbiota remains largely unexplored.

Here, we demonstrate that plant-associated *Streptomyces* sp. ATMOS43 employs a yet uncharacterized mechanism of FA detoxification through mono- and di-amino acid conjugation, thereby alleviating toxicity towards both bacteria and plants. Furthermore, we show that this activity is widespread among *Streptomyces* species. Together, our findings identify amino acid conjugation as a microbial detoxification strategy that can contribute to protection of plants under siege of plant pathogenic *Fusarium*.

## MATERIALS AND METHODS

### Microbial and plant growth conditions and extraction of metabolites

The plant used in the studies as *Arabidopsis thaliana* Columbia accession (Col-0). Seeds were surface-sterilized and stratified in sterile water at 4 °C for 48 h prior to sowing on ½-strength MS agar media. 20 seedlings of two-week old plants were transferred to 100 mL shake flasks containing 15 mL ½ MS media. Plants were infected with 5X10^5^ CFU of *Fusarium oxysporum f. sp. radicis lycopersici* (*Fox*) spore stock [30]. Sterile plants and *Fusarium* cultures were prepared the same way. The flasks were incubated for three days while shaking at 60 rpm in a climate-controlled plant growth chamber at 21 °C, 50% relative humidity, under 16 h light/8 h dark cycles. All cultures were filtered using 0.22 µM filters and 1 g of Diaion HP20 resin was added to the filtrate and incubated shaking for two hours. The HP20 was filtered and washed with Mili-Q water and then extracted with MeOH three times by soaking overnight. Extracts were dried under vacuum and stocks of 20 mg/mL were prepared in MeOH.

*S. coelicolor* A3(2) M145 was obtained from the John Innes Centre strain collection in Norwich, UK; *Streptomyces* sp. ATMOS43 was obtained from a field experiment with *Arabidopsis thaliana* ecotype Mossel (Msl) at the Mossel area at the ‘Hoge Veluwe’ in the Netherlands [31]. Routine cultivation and genetic manipulation of *Streptomyces* was done according to protocols described in the *Streptomyces* laboratory manual [32].

To screen the ATMOS collection, minimal media (MM) agar [32, 33] with 1% mannitol and 1% glycerol (w/v) as carbon sources was supplemented with crude extracts of fungal, plant or fungal-plant cultures at a final concentration of 50 µg/mL. ATMOS isolates were spotted by inoculating 3 µL of spore stocks and incubated for five days. Subsequently growth was evaluated and antifungal activity was assessed by inoculating 7X10^4^ CFU of *Fox* spore stock in the centre of the petri dish and incubated for additional three days.

### LC-MS based metabolite profiling, comparative metabolomics, and molecular networking

Metabolome analysis was performed as described in Supplementary Materials and Methods. Briefly, *Streptomyces* sp. ATMOS43 was cultured with and without *Fox*-infected plant extracts (50 µg/mL) or fusaric acid (Sigma-Aldrich, CAS #536-69-6) treatments by inoculating 1 X 10^7^ CFU in 10 mL ISP2 media and incubated at 30 °C. Fusaric acid concentration, duration of incubation, and time-point of treatment varied according to the described experiment. For LC-MS/MS analysis, HP-20 resin was added to the cultures at 1 g per 15 mL culture and incubated shaking for 3 h. Resin was collected by filtration, washed with Milli-Q water, and extracted with MeOH three times by soaking overnight. Extracts were dried under vacuum and stocks of 20 mg/mL were prepared in MeOH. LC-MS/MS data were deposited in the MassIVE public data repository (MSV000102610).

Raw data obtained from LC-MS analysis were converted to mzXML centroid files using Shimadzu LabSolutions Postrun Analysis. The files were imported into Mzmine 2.35 for data processing [34]. Methods and parameters used for data processing in Mzmine, statistical analysis, and molecular networking are described in Supplementary Materials and Methods.

For FA levels measurements in cultures of ATMOS strains were carried out with multiple reaction monitoring (MRM) as described in Supplementary Materials and Methods.

### Synthesis of the conjugate fusaric acid-serine (FA-Ser)

FA-Ser was synthesized using the N,N-dicyclohexylcarbodiimide method used by K. Veera Reddy et al., for the synthesis of picolinyl amino acids with modification [35]. Briefly, equimolar amounts of FA (Sigma-Aldrich, CAS #536-69-6), L-serine ethyl ester hydrochloride (Sigma-Aldrich, CAS # 26348-61-8), triethylamine (Sigma-Aldrich, CAS #121-44-8), and 1.05 equiv. N,N′-dicyclohexylcarbodiimide (Sigma-Aldrich, CAS #538-75-0) were dissolved in CH_3_CN:THF (2:1) and stirred at room temperature for three days. The crude ester was purified by silica gel column chromatography and EtOAc:hexane elution. Hydrolysis of the ethyl ester was achieved by suspending the product in milli-Q water and dropwise addition of NaOH over one hour with stirring and heating at 80 °C. The product was crystalized by the addition of 1 equiv. HCl. Further purification was achieved by HPLC using Waters preparative system comprised of 1525 pump, 2707 autosampler, and 2998 PDA detector. The product was injected into a SunFire C_18_ column (10 μm, 100 Å, 19 × 150 mm). The column was run at a flow rate of 12.0 mL/min, using H_2_O for solvent A and MeOH for solvent B. The gradient used was 20%-100% B for 30 min and 100% B for 5 min with FA-Ser eluting at 12 min. Final yield was 750 mg starting with 1 g FA.

### Co-culture of Streptomyces sp. ATMOS43 and *S. coelicolor* M145

*Streptomyces* sp. ATMOS43 and *S. coelicolor* M145 were inoculated in ISP2 and incubated for 24 h (early stationary phase). 500 µL of each culture was inoculated in fresh 10 mL ISP2 with and without 100 µM FA and incubated at 30 °C for four days. Cultures were centrifuged, the supernatant was alkalinized with 1 N KOH, and total actinorhodin levels were determined by measuring absorbance at 640 nm.

### FA and FA-Ser toxicity to Arabidopsis seedlings

Seeds were surface sterilised using 40% bleach, washed thoroughly with sterile water and stratified in sterile water at 4 °C for 48 h before sowing on ½ MS agar medium. Twelve 1-week-old seedlings were transferred to ½ MS medium supplemented with 60 µM FA, 60 µM FA-Ser or DMSO for solvent control and incubated for an additional week. Fresh shoot biomass of individual seedlings was measured immediately after harvest.

### Genome sequencing, assembly, and annotation of *Streptomyces* sp. ATMOS43

*Streptomyces* sp. ATMOS43 was grown in TSBS supplemented with 10mM MgCl2 and incubated with 200 rpm shaking at 30 °C. Genomic DNA was isolated by phenol-chloroform extraction as described previously [46]. Sequencing was outsourced to Novogene Co., Ltd (UK). PCR-free library preparation was performed to avoid sequencing biases. The sequencing was performed on Illumina platform in paired end mode with a read-length of 250 bp. Raw data with adapter sequences or low-quality sequences were filtered using SOAPnuke [55] to produce clean reads. Clean reads were then assembled using Shovill (v1.1.0) with SPAdes (v3.14.1) [56]. Genome annotation was performed using Prokka 1.14.6 [36]. This Whole Genome Shotgun project has been deposited at DDBJ/ENA/GenBank under the accession JCBBYW000000000. The version described in this paper is version JCBBYW010000000.

### Generation of phylogenetic trees

For the MBT collection, the phylogenetic tree was generated using 16S rRNA gene sequences extracted from the available full genome sequences. For the ATMOS collection, the phylogenetic tree was generated using 16S rRNA gene sequences described by van der Meij, et al. [31]. The sequences were aligned and the tree was generated using Geneious Alignment and Geneious Tree Builder in geneious prime (2025.0.2) and the Tamura-Nei genetic distance model. Phylogenetic trees were visualized and annotated using iTOL v7 [37].

### Proteomics sample preparation and analysis

We applied proteomining to analyse the differential expression of proteins in response to FA or FA-Ser treatments [38–40]. 10X10^7^ CFU of *Streptomyces* sp. ATMOS43 were inoculated in 10 mL ISP2 media in four replicates and incubated for 24 h at 30 °C. Cultures were treated with 100 µM FA or FA-Ser (DMSO was used as solvent control) and incubated for additional four hours. Sample preparation and proteomics analysis were performed as described in Supplementary Materials and Methods. Detailed parameters used for analysis were submitted together with raw data. The mass spectrometry proteomics data have been deposited to the ProteomeXchange Consortium via the PRIDE [41] partner repository with the dataset identifier PXD082028.

### Gene constructs and cloning

*Escherichia coli* strain DH5α was used for routine cloning and plasmid propagation. *E. coli* ET12567(pUZ8002) [42] was used for intergenic transfer of plasmids from *E. coli* to *Streptomyces*. *E. coli* strains were grown in Luria-Bertani (LB) medium at 37 °C. The parental strain for all mutants was *S. coelicolor* A3(2) M145 [43]. *Streptomyces* strains were grown on soya flour mannitol (SFM) for conjugation and sporulation at 30 °C. Media were supplemented with the appropriate antibiotics when required (apramycin, 50 µg/mL; ampicillin, 100 µg/mL; kanamycin, 50 µg/mL; chloramphenicol, 25 µg/mL; nalidixic acid, 10 µg/mL; thiostrepton, 20 µg/mL).

Primers used in this work are listed in **Table S2**. DNA used for cloning was amplified with Q5 polymerase (New England Biolabs) following manufacturer’s instructions. Constructs were assembled using NEBuilder HiFi DNA Assembly Master Mix (New England Biolabs).

The unstable multicopy vector pWHM3-oriT [44] was used to generate a knockout mutant of SCO0436 in *S. coelicolor* A3(2) M145, using a method described previously [45]. We replaced nucleotide positions +1 to +122 relative to the translational start site of SCO0436 with an apramycin resistance cassette. For this, approximately 1500 bp up- and downstream regions of SCO0436, with the *loxP*-*aac3(IV)-loxP* cassette in between, were cloned into pWHM3-oriT. Subsequent conjugation of pRS099 into *S. coelicolor* M145 resulted in strain mRS018 (ΔSCO0436::Apra). The apramycin resistance cassette was subsequently removed using the Cre-expressing construct pUWL-Cre [46], yielding the clean knockout strain mRS019 (ΔSCO0436).

For expression of *rpmF* from *Streptomyces* sp. ATMOS43 in *S. coelicolor* M145, the open reading frames of the three *rpmF* paralogues were amplified from the genome of ATMOS43 (*rpmF1*, ATMOS43_TMLOC00453; *rpmF2*, ATMOS43_TMLOC00126; and *rpmF3*, ATMOS43_TMLOC04367). The*ermE* promoter was amplified from pHM10a [47]. The three rpmF genes were cloned under expression of the *ermE* promoter into pSET152 digested with EcoRI and BamHI, yielding expression constructs pRS104, pRS105 and pRS106, respectively (**Table S3**). The constructs were conjugated into strain ΔSCO0436, resulting in strains ΔSCO0436::P*ermE-rpmF1*, ΔSCO0436::P*ermE-rpmF2*, and ΔSCO0436::P*ermE-rpmF3*,

### Metal ion supplementation and PAR assay

To evaluate how metal ion supplementation influences FA toxicity in *Streptomyces* sp. ATMOS43 and *Streptomyces coelicolor* M145, 5X10^6^ CFU from each spore stock were inoculated into 5 mL ISP2 medium, with or without 100 µM FA, in quadruplicate. The R5 trace element solution was added to achieve final concentrations of 1.7 µM Zn²⁺, 4.4 µM Fe³⁺, 0.2 µM Cu²⁺, and 0.6 µM Mn²⁺. For single-metal supplementation, CuSO₄, FeCl₃ or ZnCl₂ was added at concentrations matching those supplied by the trace element solution. At each time point, cultures were centrifuged, and the resulting biomass pellets were washed with Milli-Q water. The biomass was then dried at 60 °C for 48 h before determining dry mass.

4-(2-Pyridylazo)resorcinol (PAR) (Sigma-Aldrich, CAS 1141-59-9) was used to evaluate the relative Zn²⁺- chelation activity of FA and FA-Ser, following a previously described method with minor modifications [48]. A 100 µM PAR working solution was prepared in buffer by dissolving 2.12 g of HEPES [4-(2- hydroxyethyl)-1-piperazineethanesulfonic acid] in 40 mL Milli-Q water and adjusting the pH to 8 with 5 M NaOH. Stock solutions consisted of 500 µM ZnCl₂ in Milli-Q water and 20 mM FA or FA-Ser in DMSO. For each reaction, 120 µL mixtures were prepared by combining 60 µL PAR working solution with 60 µL sample solution, giving final concentrations of 50 µM PAR, 50 µM ZnCl₂, and 100, 250 or 500 µM FA or FA-Ser. Ethylenediaminetetraacetic acid (EDTA; 100 µM) was used as positive control. Total absorbance was measured at 490 nm.

## RESULTS

### Growth responses of root-endophytic *Streptomyces* to *Fusarium*-associated toxins

To investigate how root-associated streptomycetes respond to plant and pathogen-derived metabolites, we established a controlled bioassay in which *Arabidopsis thaliana* seedlings were infected with *Fusarium oxysporum* f. sp. *radicis lycopersici* (*Fox*) in culture flasks (**Fig. 1a**). After 3 days of incubation, plant disease symptoms and *Fox* colonization of roots were evident. Spent media from this co-culture—together with media blank, *Arabidopsis,* and *Fox* monoculture controls—were extracted with methanol and added to minimal medium (MM) agar supplemented with 1% mannitol and 1% glycerol. Next, *Streptomyces* strains from our in-house collection of *Arabidopsis* root endophytes (ATMOS collection) [31] were inoculated from spore stocks. After 5 days of incubation, several strains displayed impaired colony growth in the presence of the *Fox* extract and even more pronounced growth impairment by the *Fox*+*Arabidopsis* co-culture extract relative to control conditions (**Fig. 1b**). Colonies were subsequently tested for antifungal activity by inoculating *Fox* in the plate centre; after 3 days of incubation, *Streptomyces* sp. ATMOS43 grew robustly in all conditions and retained antifungal activity against *Fox* (**Fig. 1c**). This suggests that *Streptomyces* sp. ATMOS43 can sustain growth and antifungal activity in the presence of metabolites produced in the *Fox* monoculture and in the co-culture of *Fox*+*Arabidopsis,* whereas growth of other isolates was suppressed.

**Fig 1.**
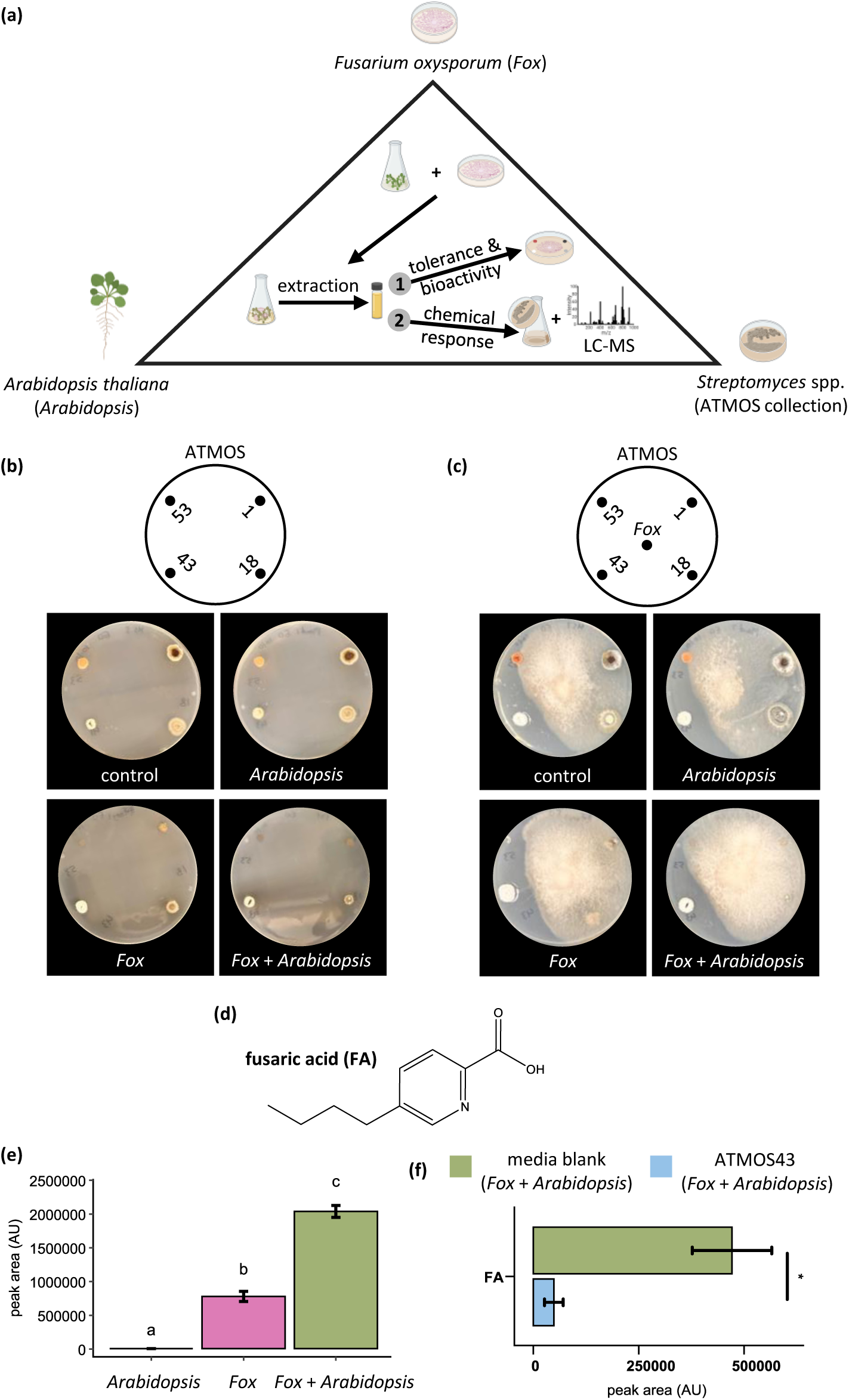
Experimental design and tolerance of *Streptomyces* sp. ATMOS43 to toxins produced by *Fusarium oxysporum* (*Fox*). **a)** Experimental setup used in this work. Arabidopsis seedlings were infected with *Fox* spores and incubated for three days. Culture supernatant was extracted and added to MM agar plates for the inoculation of ATMOS strains or analysed using LC-MS. **b)** Growth of four ATMOS strains on MM agar plates in the presence or absence of *Arabidopsis*, *Fox*, or *Fox*+*Arabidopsis* culture extract. **c)** Antifungal bioassay of ATMOS strains against *Fox* in the presence or absence of *Arabidopsis*, *Fox*, or *Fox*+*Arabidopsis* culture extract. Fungal growth is consistently inhibited by *Streptomyces* sp. ATMOS43 in all conditions. **d)** Chemical structure of FA. **e)** FA levels in *Arabidopsis*, *Fox*, and *Fox*+*Arabidopsis* cell-free culture supernatant. Levels indicate increased FA production in *Fo*x+*Arabidopsis* co-culture relative to *Fox* monoculture. Different letters show statistically significant differences (one-way ANOVA followed by Tukey’s HSD test, *p*-value < 0.05). **f)** Residual FA levels in *Streptomyces* sp. ATMOS43 in cultures supplemented with crude extract of *Arabidopsis*+*Fox* culture extract compared to media control with the same treatment. Two-sample unpaired Student’s t-test showing the significant differential levels. Asterisks indicate statistically significant differences (*p*-value < 0.05). Error bars represent standard error (n=3). AU: arbitrary units.

To identify *Fox* metabolites produced under these experimental conditions, we analysed extracts from *Fox* and *Fox*+*Arabidopsis* cultures using Liquid Chromatography-Mass Spectrometry (LC-MS). Among the known *Fox*-derived mycotoxins, which were previously identified using LC-MS [49], FA was the only one detected in our cultures, indicating that FA is the dominant toxin in this bioassay. Notably, our analysis revealed a marked increase in FA levels in *Fox*+*Arabidopsis* co-culture relative to the *Fox* monoculture (**Fig. 1d**, **1e**, and **S1**). This is in line with the observed higher toxic effect of the co-culture extract on growth of the tested bacterial strains (**Fig. 1b**). To assess whether tolerance to FA correlates with the growth inhibition observed in the initial screen, we determined the minimum inhibitory concentration (MIC) of FA across the previously tested ATMOS strains. After one week of incubation, *Streptomyces* sp. ATMOS43 displayed a higher MIC than the other strains (**Fig. S2**).

*Streptomyces* sp. ATMOS43 was then cultured in ISP2 medium under the same treatment conditions described above, with corresponding blank medium controls. We observed a significant depletion of FA derived from *Fox*+*Arabidopsis* extracts as compared to the media control (**Fig. 1f**). Notably, FA was not detected in *Streptomyces* sp. ATMOS43 cultures or the corresponding blank medium control supplemented with the *Fox* monoculture extract. Next, we performed a direct feeding experiment where *Streptomyces* sp. ATMOS43 was cultured in media with 100 µM pure FA (**Fig. S3a**). The residual levels of FA were significantly lower in *Streptomyces* sp. ATMOS43 cultures compared to media blank supplemented with the same FA concentration (**Fig. S3b**). We similarly performed the FA feeding experiment on the other ATMOS strains included in the initial screening (**Fig. 1b**). FA inhibited the growth of *Streptomyces* sp. ATMOS1 and *Streptomyces* sp. ATMO53 (**Fig. S4a**). *Streptomyces* sp. ATMOS18 continued to grow under FA treatment, however, FA levels in culture extracts remained similar to the levels in media blanks treated with the same FA concentration (**Fig. S4b**). Together, these results suggest that the growth and antifungal activity of *Streptomyces* sp. ATMOS43 in the presence of *Fox*-derived culture extracts are associated with specific mechanism(s) to metabolize or detoxify FA.

### Amino acid conjugation as a new mechanism of FA detoxification by *Streptomyces* sp. ATMOS43

*Streptomyces* species adopt various detoxification mechanisms to tolerate toxins and other natural products in the environment [50–52]. To determine whether the observed FA tolerance reflects a specific detoxification mechanism, we investigated potential transformation or degradation of FA by *Streptomyces sp.* ATMOS43. For that, we subjected our LC-MS data to Feature-Based Molecular Networking (FBMN) analysis using the Global Natural Products Social Molecular Networking (GNPS) web platform [53]. This approach enables relative quantification of mass features across different conditions, and clusters them based on MS/MS fragmentation similarities. Additionally, we used MS2LDA, which identifies molecular ‘fingerprints’ in MS/MS spectra and assigns these into Mass2Motifs [54]. The MS/MS fragmentation spectrum of pure FA was used to identify its characteristic Mass2Motif (**Fig. S5** and **S5b**), and the corresponding mass features were mapped onto the molecular network. Two molecular families were assigned as FA-related and were detected only when *Fox*+*Arabidopsis* extract was added to cultures of *Streptomyces* sp. ATMOS43 **(Fig. 2a)**. To confirm that these features are FA-related, we analysed the LC-MS results of *Streptomyces* sp. ATMOS43 cultures in the presence of pure 100 µM FA. Five out of eight of the FA-related mass features were identified by MS2LDA (**Fig. S5c**). Analysis of the corresponding MS/MS spectra indicated amino acid conjugation of FA, with level 3 metabolite identification achieved (**Fig. S6**) [55]. We identified both single amino acid conjugates (SACs) and dipeptidyl conjugates (DPCs) of FA. An assortment of canonical amino acids, including alanine, serine, threonine, and glutamine, were conjugated to FA via amide bond formation; DPCs consisted of variable first amino acids, whereas the second amino acid was consistently glutamine (**Fig. 2b** and **S6**). To confirm the identity of the annotated FA conjugates, we chemically synthesized FA-Ser—the primary conjugate detected—and verified it by LC-MS as well as 1D and 2D NMR (**Table S1** and **Fig. S7-S15**). We found that the LC-MS peak of the synthesized compound matched the peak in the *Streptomyces* sp. ATMOS43 culture extract in terms of retention time, exact mass, and MS/MS spectrum (**Fig. S16**).

**Fig 2.**
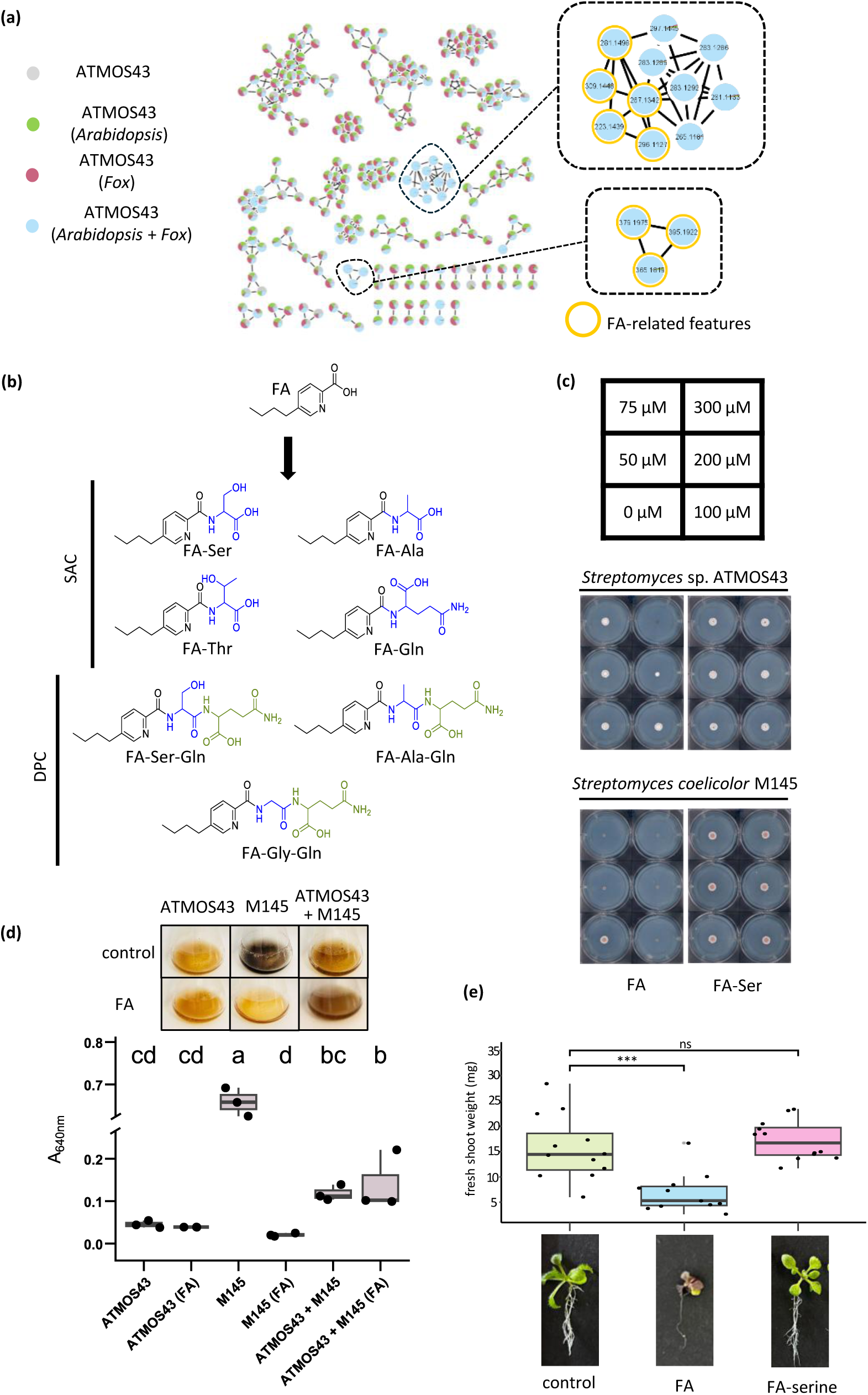
Identification of amino acid conjugation as a mechanism of FA detoxification by *Streptomyces* sp. ATMOS43. **a)** Feature-based molecular network (FBMN) of the ions detected in the crude extracts of *Streptomyces* sp. ATMOS43 cultures under the different treatment conditions. Magnified molecular families include features with FA Mass2Motif as identified by MS2LDA (yellow node borders). **b)** Chemical Structures of single amino acid conjugates (SAC) and di-peptidyl conjugates (DPC) identified in this work. **c)** Minimum inhibitory concentration (MIC) assay of FA and FA-Ser against *Streptomyces* sp. ATMOS43 and *S. coelicolor* M145. Increased resistance to FA is observed for *Streptomyces* sp. ATMOS43 compared to *S. coelicolor* M145 while no toxicity of FA-Ser is evident for both strains at all tested concentrations. **d)** Box and whisker plot of total UV absorbance at 640 nm measured for the supernatant of 4-days old cultures showing the significant increase of actinorhodin levels in co-culture of *S. coelicolor* M145 with *Streptomyces* sp. ATMOS43 compared to the monoculture in the presence of 100 µM FA. One-way ANOVA followed by Tukey’s HSD test (n=3). Different letters show statistically significant differences (*p*-value < 0.05). **e)** Comparison of FA and FA-Ser toxicity to *Arabidopsis* seedlings. Box and whisker plot represent fresh shoot biomass measurements. FA treatment results in significant growth inhibition of *Arabidopsis* plants while no detectable toxicity is observed for FA-Ser. One-way ANOVA followed by Tukey’s HSD test (n=12). ***; *p*-value < 0.001, ns; non-significant.

We next tested if amino acid conjugation affects FA toxicity. For this, we compared the toxicity of the chemically synthesized FA-Ser to that of FA on *Streptomyces* sp. ATMOS43 and the model strain *Streptomyces coelicolor* M145. We performed a MIC assay using a concentration gradient of FA and FA-Ser. FA exhibited adverse effects on colony growth at 50 µM in *S. coelicolor* M145 and at 200 µM in *Streptomyces* sp. ATMOS43 (**Fig. 2c**). In contrast, FA-Ser displayed no detectable adverse effect on colony growth of either two strains at concentrations up to 300 µM. Notably, *S. coelicolor* showed no evidence of FA detoxification or degradation in LC-MS measurements. Similarly, FA-Ser did not show detectable toxicity on growth of the other ATMOS strains tested in the initial screening (**Fig. S17**). Subsequently, we tested if FA detoxification by *Streptomyces* sp. ATMOS43 would mitigate its toxicity towards *S. coelicolor*, whose growth can be visually assessed through the production of the characteristic blue pigment actinorhodin [56]. For this, each strain was inoculated with or without 100 µM FA in mono- or coculture in ISP2 medium and incubated for four days. Levels of actinorhodin produced by *S. coelicolor* were measured by total UV absorbance at 640 nm as previously described [32]. In the presence of FA, actinorhodin production was partially restored when *S. coelicolor* was co-cultured with *Streptomyces* sp. ATMOS43, indicating alleviation of FA toxicity in the co-culture setting (**Fig. 2d**).

Additionally, we tested if amino acid conjugation would mitigate the toxicity of FA to plants. We transferred 1 week old *Arabidopsis* seedlings to ½ MS media supplemented (or not) with 60 µM FA or FA-Ser and determined shoot biomass after an additional week of incubation. FA significantly inhibited *Arabidopsis* seedling growth, whereas FA-Ser had no detectable adverse effects **(Fig. 2e)**. Together, these results indicate that amino acid conjugation contributes to FA resistance in *Streptomyces* sp. ATMOS43 and can effectively diminish FA toxicity towards *Arabidopsis* and FA-sensitive *Streptomyces*.

### Prevalence of FA conjugation in streptomycetes

To investigate how prevalent amino acid conjugation to FA is in Streptomycetaceae and how widespread is the formation of dipeptidyl conjugates (DPCs) of FA, we screened our in-house actinobacterial collection of *Streptomyces* and *Kitasatospora* strains isolated from soil (MBT collection; [57]) and compared their response to FA to the collection of *Streptomyces* strains isolated from *Arabidopsis* roots (ATMOS collection; [31]). All strains were inoculated from spore stocks in ISP2 media, incubated for 24 h before treatment with 100 µM FA and then incubated for an additional 24 h in presence of FA. Cultures were extracted and analysed by LC-MS as described above. The results showed that several strains can form single amino acid conjugates (SACs) while others form both SACs and DPCs. Out of 68 MBT strains, 14 could conjugate amino acids to FA from which DPC were formed by 3 strains (**Fig. 3a)**. From the 30 ATMOS strains, 13 could conjugate FA from which DPC were detected in 5 strains. Subsequently, we mapped FA conjugation activity on a phylogenetic tree with the strains of each collection. For this, we constructed separate phylogenetic trees of the two collections based on 16S rRNA sequences. The results show that amino acid conjugation of FA is broadly distributed across phylogenetically distant *Streptomyces* strains (**Fig. S18**). Collectively, these results revealed that amino acid conjugation is prevalent among actinobacterial strains from soil and plant-associated environments.

**Fig 3.**
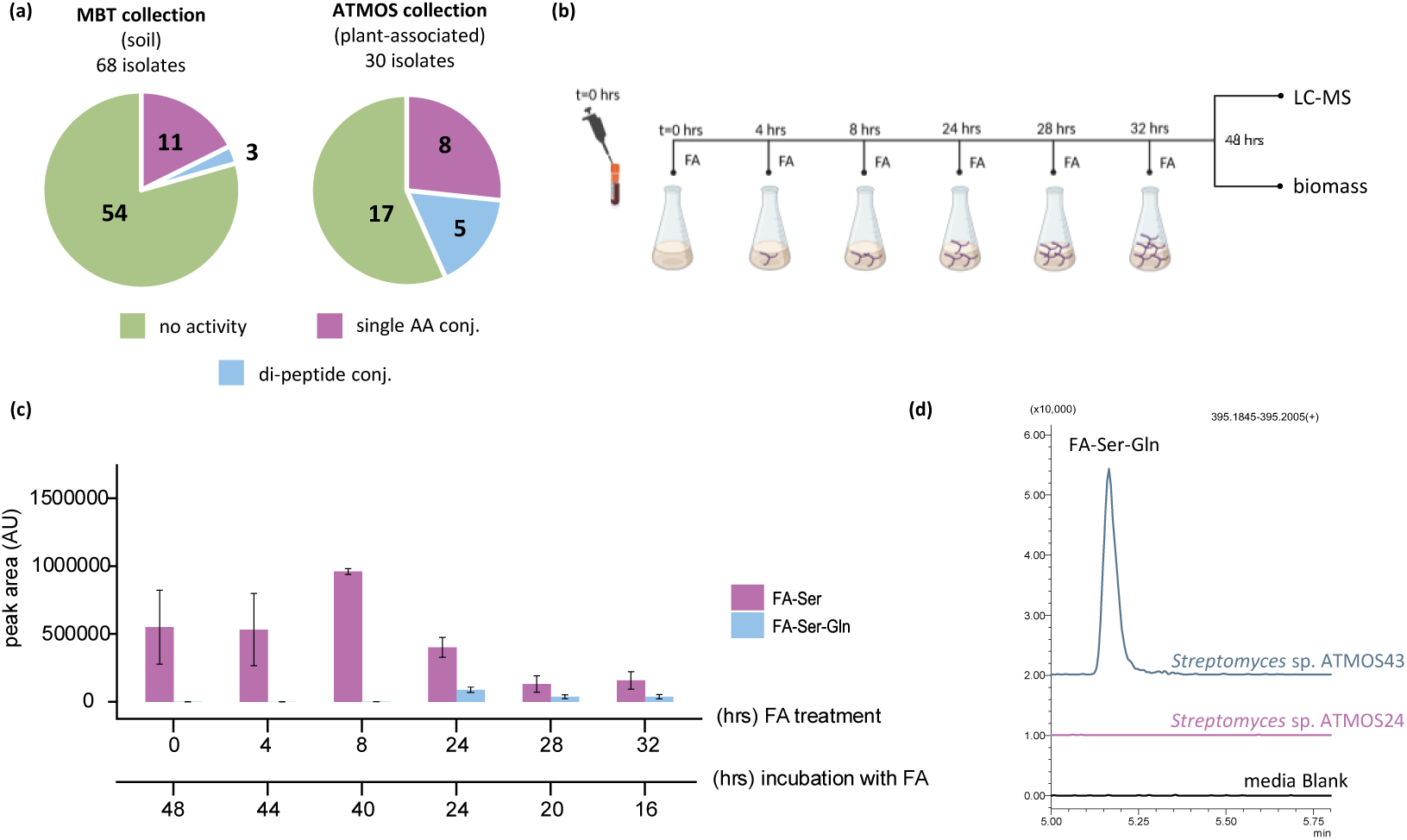
Prevalence of FA aminoacylation activity among *Streptomyces* strains and the sequential amino acid conjugation of FA by *Streptomyces* sp. ATMOS43. a) Pie charts illustrate the prevalence of SAC and DPC formation activity in soil-dwelling (MBT collection) and plant-associated (ATMOS collection) *Streptomyces* isolates. Results indicate a higher prevalence of FA conjugation activity among plant-associated isolates. **b)** Experimental setup of the time-series FA feeding to *Streptomyces* sp. ATMOS43 cultures at different stages of growth. 100 µM FA was added to a spore inoculum of *Streptomyces* sp. ATMOS43 (t = 0) or to 4, 8, 24, 28, or 32 h cultures. Three cultures of each time points were extracted for LC-MS profiling of SAC and DPC while the biomass of identical three cultures were harvested for growth assessment. **c)** Differential patterns of FA-Ser and FA-Ser-Gln formation by *Streptomyces* sp. ATMOS43 upon FA feeding at different stages of growth. FA feeding at early growth stages (0, 4, and 8 h) results in the formation of FA-Ser without detectable formation of FA-Ser-Gln. FA feeding at late growth stages (24, 28, and 32 h) results in the formation of both FA-Ser and FA-Ser-Gln. **d)** Extracted ion chromatograms of FA-Ser-Gln after FA-Ser feeding to *Streptomyces* sp. ATMOS43 and *Streptomyces* sp. ATMOS24. FA-Ser-Gln formation is only detected for *Streptomyces* sp. ATMOS43 indicative of sequential conjugation of amino acids to FA.

### Sequential amino acid conjugation of FA by *Streptomyces* sp. ATMOS43

Given that the detoxification of FA via aminoacylation has not been described previously, we sought to investigate the amino acid conjugation dynamics. We designed a set-up where 100 µM FA was fed to *Streptomyces* sp. ATMOS43 cultures at different stages of growth (**Fig. 3b**). FA was added to growing cultures at 0, 4, and 8 h corresponding to lag and early exponential phases and 24, 28, and 32 h corresponding to late exponential and stationary phases of growth (**Fig. S19**). All samples were analysed after 48 h of incubation via LC-MS to investigate the aminoacylation pattern. We specifically monitored the formation of FA-Ser and FA-Ser-Gln as the primary SACs and DPCs, respectively. In parallel, similar cultures were set for biomass measurements to compare growth to untreated (control) cultures. Results indicate that FA feeding to cultures in the lag phase results only in SACs (**Fig. 3c**), while the biomass of the cultures is significantly lower than in control conditions without FA (**Fig. S20**). The formation of both SACs and DPCs is observed for FA treatment of cultures in late exponential or stationary growth phases (**Fig. 3c**). Biomass yields were comparable to untreated controls, while FA addition at 32 h even yielded significantly higher biomass (**Fig. S20**). These results indicate that amino acid conjugation to FA is a sequential mechanism where one amino acid is first bound to FA and subsequently Gln is conjugated to the C-terminal of the peptide. To test this, we performed a feeding experiment with 100 µM FA-Ser added to 24 h cultures. LC-MS results showed FA-Ser-Gln formation when cultures were extracted after an additional 24 h of incubation (**Fig. 3d**). Moreover, we performed the FA-Ser feeding with *Streptomyces* sp. ATMOS24, which is one of the strains tested in our previous screening and found to only form SAC, including FA-Ser, when treated with FA (**Fig. 3a**). *Streptomyces* sp. ATMOS24 failed to conjugate a second amino acid to FA-Ser (**Fig. 3d**). Collectively, these results indicate that FA aminoacylation proceeds sequentially, with SAC formation preceding addition of Gln to generate DPCs. The inability of *Streptomyces* sp. ATMOS24 to convert FA-Ser into FA-Ser-Gln further suggests that DPC formation requires an additional metabolic conversion that is absent or inactive in this strain and likely in the other strains for which only SACs were detected upon FA treatment (**Fig. 3a**).

### FA toxicity involves disruption of zinc homeostasis

The toxic effects of FA are well documented for plants [25–27], but the mode of action for microorganisms is not known. To investigate how FA affects growth of streptomycetes, we compared the proteomic response of *Streptomyces* sp. ATMOS43 to FA and its conjugate FA-Ser. *Streptomyces* sp. ATMOS43 was grown in ISP2 media for 24 h, supplemented with 100 µM FA or FA-Ser and grown for an additional 4 h. Samples were collected and analysed by quantitative LC-MS-based proteomics. FA induced a stronger proteomic response than FA-Ser, with 16 differentially abundant proteins (DAPs) detected in FA-treated cultures compared with 5 DAPs in FA-Ser-treated cultures (adjusted *p*-value < 0.05, |log₂ fold change| > 1) (**Fig. 4a**). The FA-responsive proteins included high-affinity zinc uptake proteins, a cadmium/cobalt/zinc antiporter, a manganese transport protein, RpmF2/L32, EF-Tu and several metabolic or hypothetical proteins (**Fig. 4b**). These changes suggested that FA exposure causes a metal-deprivation response, with disruption of zinc homeostasis emerging as a prominent candidate mode of action of FA.

**Fig 4.**
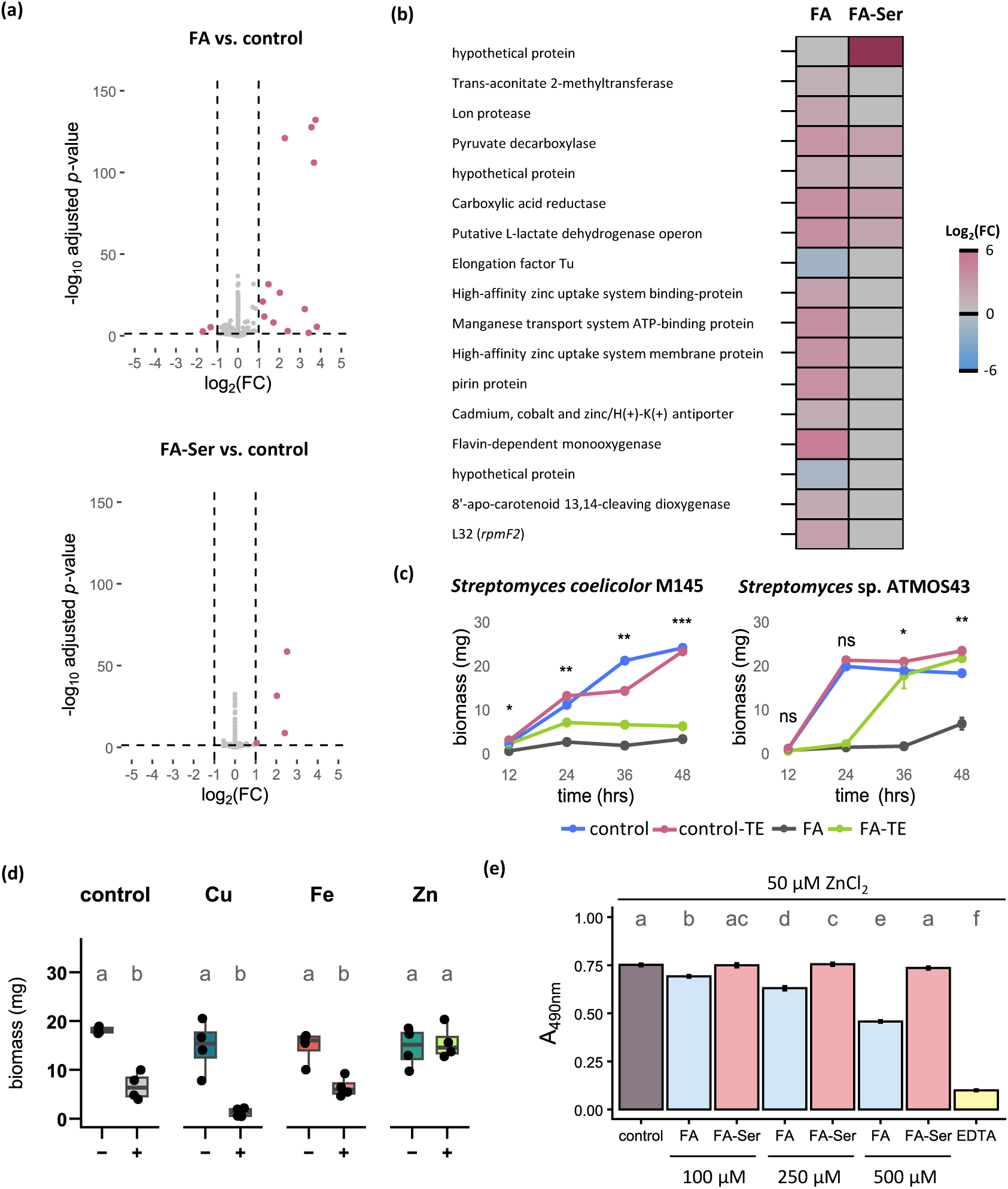
Differential effects of FA and FA-Ser on zinc homeostasis in *Streptomyces* sp. ATMOS43. **a)** Volcano plots showing LC-MS-based quantitative proteomics of *Streptomyces* sp. ATMOS43 treated with FA or FA-Ser for 4 h (n = 4). Dashed lines indicate the thresholds used for differential abundance: adjusted *p*-value < 0.05 and |log₂ fold change| > 1. **b)** Heatmap of differentially abundant proteins detected after FA or FA-Ser treatment. FA treatment affected proteins associated with metal homeostasis, translation, metabolism and stress responses, whereas FA-Ser caused a smaller proteomic response. **c)** Growth curves of *Streptomyces* sp. ATMOS43 and *S. coelicolor* M145 in the presence or absence of FA and trace elements (TE). TE supplementation partially alleviates FA toxicity in *S. coelicolor* M145 and supports recovery of ATMOS43 growth after 24 h. Statistical differences were evaluated using two sample Student’s t-test with Benjamini-Hochberg correction comparing the biomass of FA and FA-TE cultures (n=4). Asterisks indicate statistically significant differences. (*; *p*-value < 0.05, **; *p*- value < 0.01, ***; *p*-value < 0.001, ns; non-significant). **d)** Box and whisker plot showing the effect of CuSO_4_, FeCl_3_, or ZnCl_2_ supplementation on *Streptomyces* sp. ATMOS43 biomass accumulation after 48 h of growth in the presence (**+**) or absence (**-**) of FA. This shows that zinc, but not copper or iron, restores growth of *Streptomyces* sp. ATMOS43 grown in the presence of FA. One-way ANOVA, followed by Tukey’s HSD test was used to compare biomass levels (n=4). Different letters show statistically significant differences (*p*-value < 0.05). **e)** PAR assay measuring free zinc in solutions containing 50 µM ZnCl₂ and increasing concentrations of FA or FA-Ser. EDTA was included as a positive control. This shows that FA reduces free zinc in a concentration-dependent manner, whereas FA-Ser does not detectably chelate zinc. Statistical differences were evaluated by one-way ANOVA followed by Tukey’s HSD test (n = 4). Different letters indicate statistically significant differences (p < 0.05).

Because RpmF2/L32 was among the FA-responsive proteins, and L32 paralogues have previously been linked to zinc limitation in *S. coelicolor* [49], we tested whether *Streptomyces* sp. ATMOS43 RpmF proteins contribute to FA resistance. Genome analysis identified three L32 ribosomal proteins in *Streptomyces* sp. ATMOS43 while two are identified in *S. coelicolor* M145 (**Fig. S21**). Initially, we attempted to generate a knockout mutant of the L32-encoding gene *rpmF2* in *Streptomyces* sp. ATMOS43, which was upregulated following FA treatment. Following several attempts, however, *Streptomyces* sp. ATMOS43 was not accessible for genetic manipulation. Therefore, we deleted the gene encoding RpmF2 (SCO0436) from the chromosome of *S. coelicolor* and used strain M145 as a platform to study the functionality of the orthologues from *Streptomyces* sp. ATMOS43. To create the mutant ΔSCO0436 we used a method described previously. In brief, nucleotide positions +1 to +122 of SCO0436 was replaced with an apramycin resistance cassette for the generation of the knock-out construct (for plasmids and primers see **Table S2** and **S3**). The flanking regions were amplified by PCR from gDNA of *S. coelicolor* M145 and the DNA fragments were cloned into the multicopy vector pWHM3-*oriT*. The *aac*(3)*IV* apramycin resistance cassette flanked by *loxP* sites was inserted between the flanking regions. The construct was conjugated to *S. coelicolor* M145 using the methylase deficient strain *Escherichia coli* ET12567/pUZ8002 [58, 59]. Correct mutants were verified by PCR. Next, we complemented strain ΔSCO0436 with clones expressing one of the three *rpmF* paralogues from ATMOS43 using the constitutive *ermE* promoter. In this way, we generated strains ΔSCO0436::P*ermE*- *rpmF1*, ΔSCO0436::P*ermE-rpmF2*, and ΔSCO0436::P*ermE*-*rpmF3* (see Materials and Methods). The MIC assay described above with these transformants did not result in enhanced resistance of *S. coelicolor* to FA (**Fig. S22**). Thus, RpmF2 accumulation appears to be most consistent with a cellular response to FA-induced metal deprivation rather than a protective role in FA resistance.

To test whether metal availability modulates FA toxicity, we supplemented cultures with trace elements (TE) containing copper, iron and zinc. *Streptomyces* sp. ATMOS43 and *S. coelicolor* M145 were inoculated as spores in ISP2 media supplemented with TE in the presence or absence of FA, in four replicates. Growth was assessed by biomass accumulation over 48 h. For *S. coelicolor*, TE supplementation significantly improved growth in the presence of FA at all time points, although toxicity was not fully alleviated (**Fig. 4c**). In contrast, TE supplementation had little effect on *Streptomyces* sp. ATMOS43 during the first 24 h of growth. Growth recovered during the subsequent 12 h to levels comparable to the untreated controls. To determine which metal accounted for this effect, we supplemented *Streptomyces* sp. ATMOS43 cultures with CuSO₄, FeCl₃ or ZnCl₂ at equimolar concentrations present in TE. Iron and copper did not detectably alleviate FA toxicity, whereas zinc supplementation restored growth in presence of FA (**Fig. 4d**). These results indicate that FA toxicity involves disruption of zinc homeostasis.

We then tested whether FA can directly chelate zinc *in vitro* and if this activity is eliminated upon amino acid conjugation. For this, we compared the zinc-chelating activity of FA and FA-Ser using the 4-(2- pyridylazo)resorcinol (PAR) spectroscopic assay, which measures levels of free (non-chelated) zinc [48]. We prepared solutions containing 50 µM Zn and increasing concentrations of FA or FA-Ser (100, 250, and 500 µM). FA reduced free zinc levels in a concentration-dependent manner, while FA-Ser had no measurable effect across the tested concentration range (**Fig. 4e**). Finally, we showed that zinc availability did not affect the amino acid conjugation of FA by *Streptomyces* sp. ATMOS43 (**Fig. S23**), indicating that zinc does not interfere with FA detoxification by amino acid conjugation. Together, these results link FA toxicity to zinc homeostasis and show that amino acid conjugation abolishes the zinc-chelating activity of FA. The partial rescue by trace elements and the broader proteomic response suggest that additional, yet unknown cellular effects may also contribute to FA toxicity.

## DISCUSSION

Streptomycetes live in close association with plants and are exposed to similar environmental stressors as their host, including those imposed by invasions of plant pathogenic fungi. *Fusarium* species are devastating pathogens of a wide range of crops, resulting in substantial yield losses [60–62]. In this work, we investigated the potential of plant-associated streptomycetes to mitigate the toxicity of *Fusarium*-derived toxins. When plant-associated *Streptomyces* were challenged with extracts from *Fusarium*-infected *Arabidopsis* plants, we revealed a novel detoxification mechanism of FA, a key mycotoxin produced by several *Fusarium* species [24, 27]. More specifically, several *Streptomyces* strains across a phylogenetically diverse collection from soil and plant-associated environments were shown to detoxify FA via conjugation to single or double amino acids. We previously described a similar aminoacylation activity targeting the plant hormone jasmonic acid by *Streptomycetaceae* [63]. Similarly, gut-associated bacteria conjugate bile acids in the human intestine to amino acids derived from the diet [64–66]. These results suggest that aminoacylation of environmental carboxylic acid-containing metabolites is a widely distributed strategy among bacterial genera across diverse hosts and ecosystems.

Streptomycetes harbour unique biosynthetic potential and produce antifungal natural products that contribute to the suppression of *Fusarium* in plant-associated environments [67–69]. We describe a new mechanism by which streptomycetes can mitigate virulence of plant pathogenic *Fusarium*. The aminoacylation of FA reduces its toxicity towards plant seedlings and other streptomycetes, revealing a novel mechanism of toxin neutralization that is independent of the direct antifungal activity of *Streptomyces* toward *Fusarium*. This highlights the multifactorial mechanisms by which beneficial plant-associated microbes can diminish the toxic effects of pathogen-derived metabolites as well as inhibiting pathogen growth. Recently, glutamine leakage from *Arabidopsis* was identified as a major cue that drives bacterial colonization of plant roots [70]. Our work shows that *Streptomyces* sp. ATMOS43 can employ glutamine to form a second amide bond in SAC of FA; however, the added functional value for *Streptomyces* or for the plant of adding this second amino acid remains unclear and requires further investigation.

FA exerts diverse toxic effects on plant cells, including oxidative stress and mitochondrial dysfunction ultimately leading to cell death [25–27]. In cucumber plants, adding zinc or copper decreases the toxicity of FA by minimising its absorption into the plant and mitigating its toxicity likely through chelation [71]. However, FA toxicity in microbial interactions is less well understood. Using a proteomics-based approach, we identified zinc chelation as an important contributor of FA toxicity towards *Streptomyces*. FA induced a proteomic response enriched for metal-homeostasis proteins, zinc addition alleviated FA toxicity, and pure FA reduced free zinc levels *in vitro*. By contrast, FA-Ser did not detectably chelate zinc and was not toxic to *Streptomyces* nor to *Arabidopsis* growth. These findings provide a mechanistic explanation for why amino acid conjugation detoxifies FA: formation of the amide conjugate eliminates the zinc-chelating activity of the parent molecule. The upregulation of RpmF2/L32 after treatment with FA is consistent with a zinc-limitation-like response, as L32 paralogues have previously been linked to zinc starvation in *S. coelicolor* [72]. Complementation with *Streptomyces* sp. ATMOS43 RpmF paralogues did not increase FA resistance, indicating that RpmF2 accumulation is best interpreted as a marker of the cellular response to FA rather than having a role in FA resistance. In this context, we suspect that the upregulation of *rpmF2* expression in *Streptomyces* sp. ATMOS43 following FA treatment is a secondary effect of the zinc-chelating activity of FA. These observations suggest that FA exerts multi-layered toxicity, in which metal chelation is a major component, but additional cellular targets cannot be excluded.

In summary, our results show that the plant protective effects of *Streptomyces* extend beyond direct antagonism of plant pathogenic fungi and encompasses mechanisms that alleviate the toxicity of phytotoxins. Developing strategies to identify such indirect microbial bioactivities and to resolve their biosynthetic and regulatory pathways and modes of action, represent a novel avenue to uncover and harness the full functional potential of specific members of plant-associated microbiomes.

