## Supplemental Information for "Plant-associated *Streptomyces* detoxify the mycotoxin fusaric acid by amino acid conjugation"

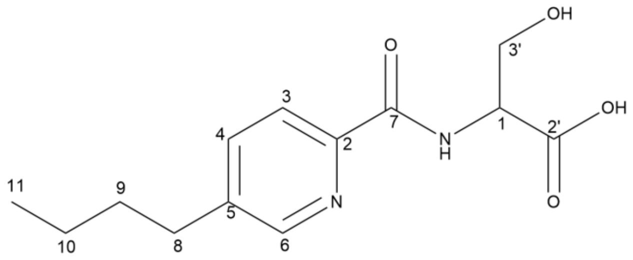


**Table S1. $^{1}H$ and $^{13}C$ NMR data of Fusaric Acid-Serine synthesized in this work (600 MHz, in DMSO-d6)**

| position | $\delta_c$, type | $\delta_H$, multiplicity, (*J* in Hz) |
|---|---|---|
| 2 | 147.2, C | |
| 3 | 121.5, CH | 7.97, d (7.9) |
| 4 | 137.5, CH | 7.86, dd (8.0, 2.2) |
| 5 | 141.3, C | |
| 6 | 148.4, CH | 8.52, d (2.2) |
| 7 | 163.5, C | |
| 8 | 31.6, $CH_2$ | 2.68, t (7.7) |
| 9 | 32.6, $CH_2$ | 1.58, m |
| 10 | 21.6, $CH_2$ | 1.30, m |
| 11 | 13.7, $CH_3$ | 0.89, t (7.4) |
| 1’ | 54.8, CH | 4.48, dt (7.8, 3.7) |
| 2’ | 171.6, C | |
| 3’ | 61.3, $CH_2$ | 3.77, dd (11.1, 3.5)<br>3.89, dd (11.1, 3.9) |
| NH | | 8.64, d (8.2) |

**Table S2. Overview of the oligonucleotides used in this work**

| Name | 5'-3' Sequence | Function |
|---|---|---|
| oRS323 | GTTGTAAAACGACGGCCAGTGGCAACATCCGGTCCCTGGAC | Amplification upstream region SCO0436 |
| oRS324 | AGTTATCCATCACCTCTAGAGGTGAGGCTCCTTTCGGACG | |
| Apra_F | TCTAGAGGTGATGGATAACTTCGT | Amplification aac3(IV) |
| Apra_R | TCTAGAGATGCGCGATAACTTC | |
| oRS333 | AGTTATCGCGCATCTCTAGAGCACCTGGTGCCCGCGTACC | Amplification downstream region SCO0436 |
| oRS330 | GCTATGACCATGATTACGCCACGACGCCATCCGGGCGGACAA | |
| oRS354 | AAGTACGACGCGGCCGCCAT | Check locus SCO0436 |
| oRS355 | CAGATCTCCGGGCGCAGGAGGC | |
| oRS239 | AACAGCTATGACATGATTACGAATTCGAGCTCGGTACCAGCCCGAC | Amplification *ermE* promoter |
| oRS262 | ATGGGTCCTCCTGTGGAGTG | |
| oRS360 | CACTCCACAGGAGGACCCATATGGCCGTACCCAAGCGCAA | Amplify rpmF2 (ATMOS43_TMLOC00126) |
| oRS359 | GGCTGCAGGTCGACTCTAGAGTCAGCCCTTCGGGTCGATCAG | |
| oRS363 | CACTCCACAGGAGGACCCATGTGGCTGTTCCGAAGCGGAAGATGTCGCG | Amplify rpmF1 (ATMOS43_TMLOC00453) |
| oRS362 | GGCTGCAGGTCGACTCTAGAGTCAGACCTCGAGGACCTGGCGC | |
| oRS366 | CACTCCACAGGAGGACCCATATGGCCGTACCCAAACGAAAGACGT | Amplify rpmF3 (ATMOS43_TMLOC04367) |
| oRS365 | GGCTGCAGGTCGACTCTAGAGTCAGCTCTCCGGGCGAAGGAG | |

**Table S3. Overview of the plasmids and constructs.**

| Name | Description | Reference |
|---|---|---|
| pWHM3-oriT | E. coli/Streptomyces shuttle vector, high copy number and unstable in *Streptomyces* | [1, 2] |
| pUWL-Cre | E. coli/Streptomyces shuttle vector expressing the Cre recombinase in *Streptomyces* | [3, 4] |
| pSET152 | E. coli/Streptomyces shuttle vector carrying *attP* site and integrase gene of ΦC31 phage for stable integration into chromosomal *attB* site of Streptomyces | [5] |
| pHM10a | E. coli/*Streptomyces* shuttle vector, harbouring the *ermE* promoter and engineered ribosome binding site | [6] |
| pRS099 | pWHM3-oriT containing SCO0436 flanking regions with apramycin resistance cassette *aac(3)IV* with *loxP* sites inserted between flanking regions | This work |
| pRS104 | pSET152 with rpmF2 (ATMOS43_TMLOC00126) under the control of *ermE* promoter | This work |
| pRS105 | pSET152 with rpmF1 (ATMOS43_TMLOC00453) under the control of *ermE* promoter | This work |
| pRS106 | pSET152 with rpmF3 (ATMOS43_TMLOC04367) under the control of *ermE* promoter | This work |

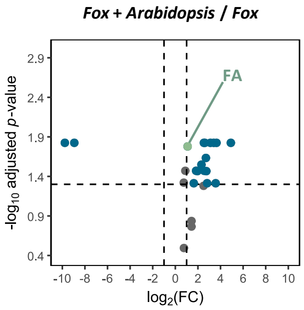


**Fig S1. Overproduction of fusaric acid (FA) by *Fusarium oxysporum* (*Fox)* upon plant colonization.** Volcano plot comparing the metabolites detected in *Fox* vs *Fox+Arabidopsis* culture extract. *Fox* overproduced metabolites include the phytotoxin FA (depicted in green). Dashed line represents the level of significance with adjusted *p*-value ≤ 0.05 and $|Log_2(FC)| > 1$.

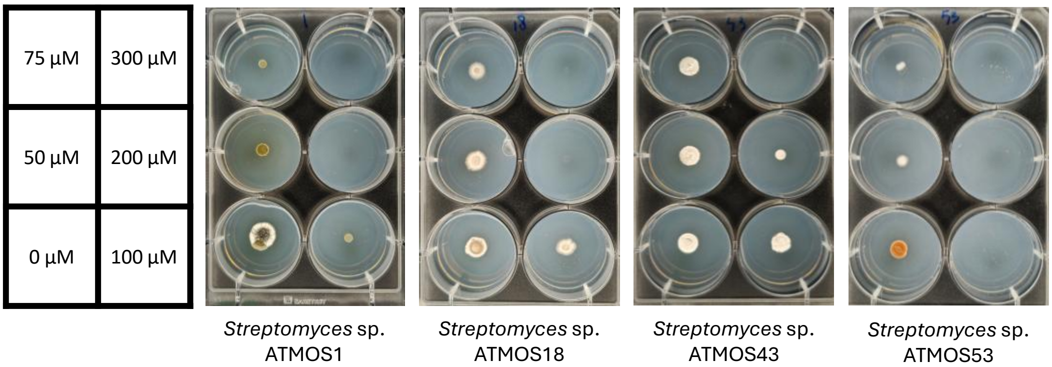


**Fig S2. Minimum Inhibitory Concentration (MIC) assay of FA across previously tested ATMOS isolates**. Streptomyces sp. ATMOS43 displayed a higher MIC than all other isolates after one week of incubation.

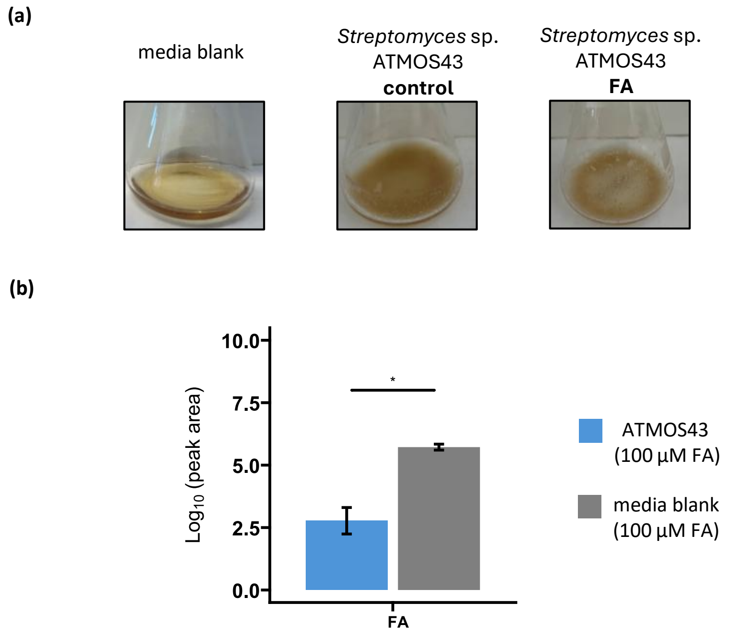


**Fig S3. Depletion of fusaric acid (FA) by *Streptomyces* sp. ATMOS43**. Residual FA levels in *Streptomyces* sp. ATMOS43 in cultures supplemented with 100 µM FA compared to media control with the same treatment. Levels indicate the depletion of FA by *Streptomyces* sp. ATMOS43. Two-sample unpaired Student's t-test showing the significant differential levels. Asterisks indicate statistically significant differences ($p$-value < 0.05). Error bars represent standard error (n=3).

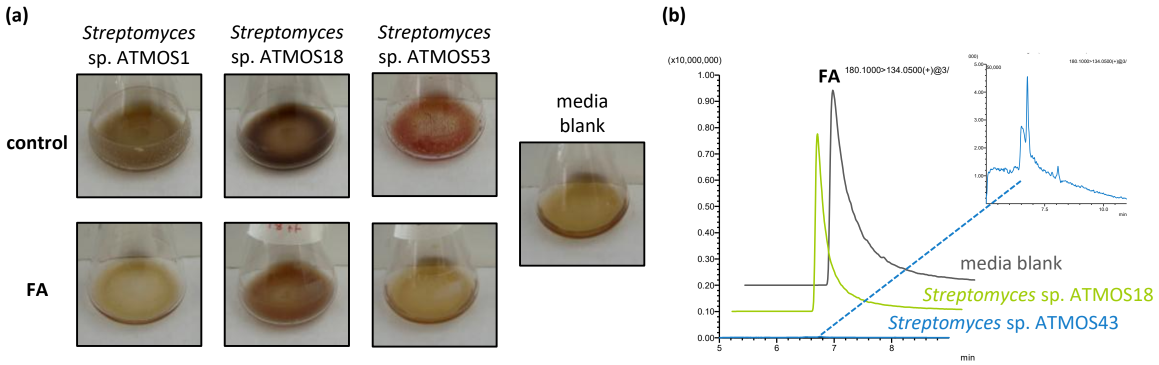


**Fig S4. Resistance and depletion of FA by ATMOS strains. a)** *Streptomyces* ATMOS strains tested in our initial screening (**Fig. 1b**) were grown in liquid ISP2 media with and without 100 µM FA. Growth was observed for *Streptomyces* sp. ATMOS18 and *Streptomyces* sp. ATMOS43 while no growth of *Streptomyces* sp. ATMOS 1 and *Streptomyces* sp. ATMOS53 was observed. **b)** Residual FA levels detected with multiple reaction monitoring (MRM) following the $m/z$ transition: 180 -> 134 in extracts of *Streptomyces* sp. ATMOS18 and *Streptomyces* sp. ATMOS43 cultures compared to media blank treated with the same concentration of FA. Levels indicate the depletion of FA by *Streptomyces* sp. ATMOS43 but not by *Streptomyces* sp. ATMOS18.

(a)

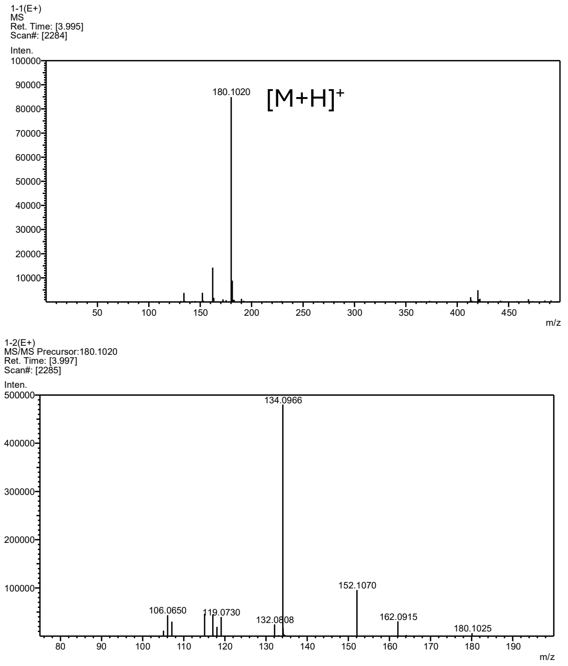


(b)

| FA motif_64_probability threshold 0.01 (1%) | | | |
|---|---|---|---|
| Feature | Probability | Feature | Probability |
| frag@134.1 | 0.2355 | loss@62.0 | 0.0181 |
| frag@162.09 | 0.1024 | loss@88.07 | 0.0175 |
| frag@152.11 | 0.069 | loss@114.02 | 0.0166 |
| frag@180.1 | 0.0645 | loss@129.05 | 0.0166 |
| frag@203.12 | 0.0374 | loss@164.03 | 0.0166 |
| loss@147.05 | 0.0321 | loss@91.03 | 0.0155 |
| frag@175.12 | 0.0274 | frag@161.11 | 0.0113 |
| frag@221.13 | 0.0269 | frag@148.11 | 0.011 |
| frag@249.12 | 0.0234 | loss@114.04 | 0.0106 |
| loss@101.05 | 0.0188 | loss@119.06 | 0.0106 |

(c)

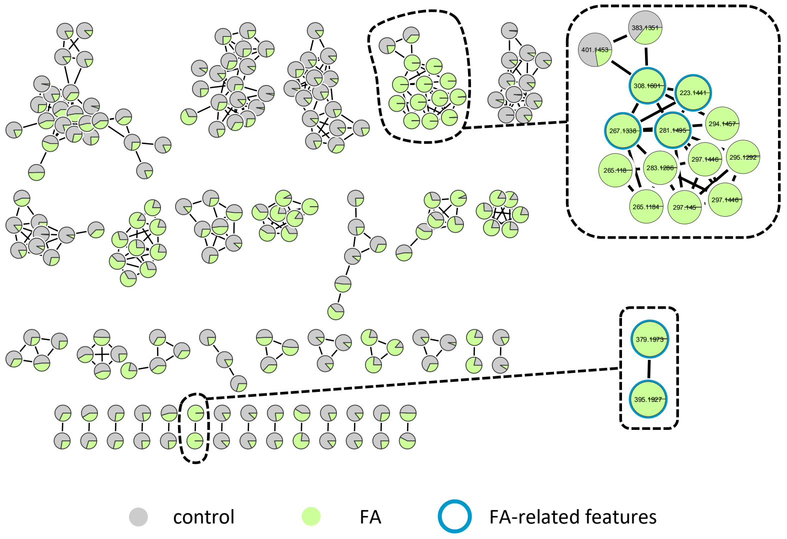


**Fig S5. Detection of fusaric acid (FA)-related features in *Streptomyces* sp. ATMOS43 culture. a)** FA (+)-HR-ESI-MS spectrum (top panel) and CID MS/MS spectrum of the $[M+H]^+$ ion (bottom panel). Spectra recorded from FA authentic standard. **b)** Mass2Motif of FA generated with MS2LDA2 (minimal feature probability 1%). **c)** Feature-based molecular network (FBMN) of the ions detected in the crude extracts of *Streptomyces* sp. ATMOS43 cultures with or without 100 µM FA treatment. Magnified molecular families include features with FA Mass2Motif as identified by MS2LDA (blue node borders).

**(a) FA-Ser**

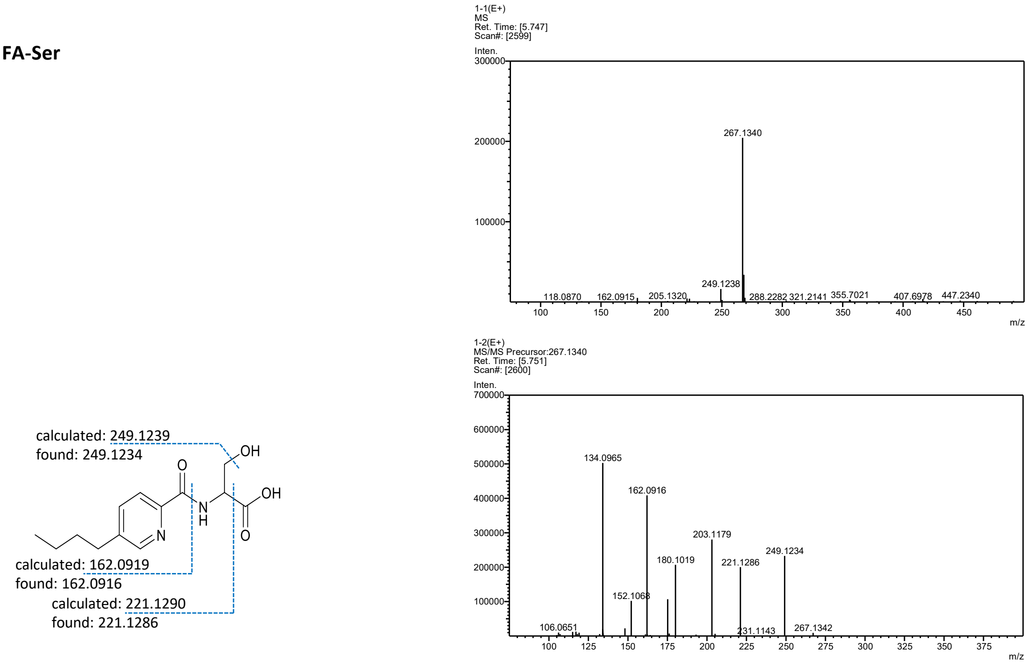


**(b) FA-Ser-Gln**

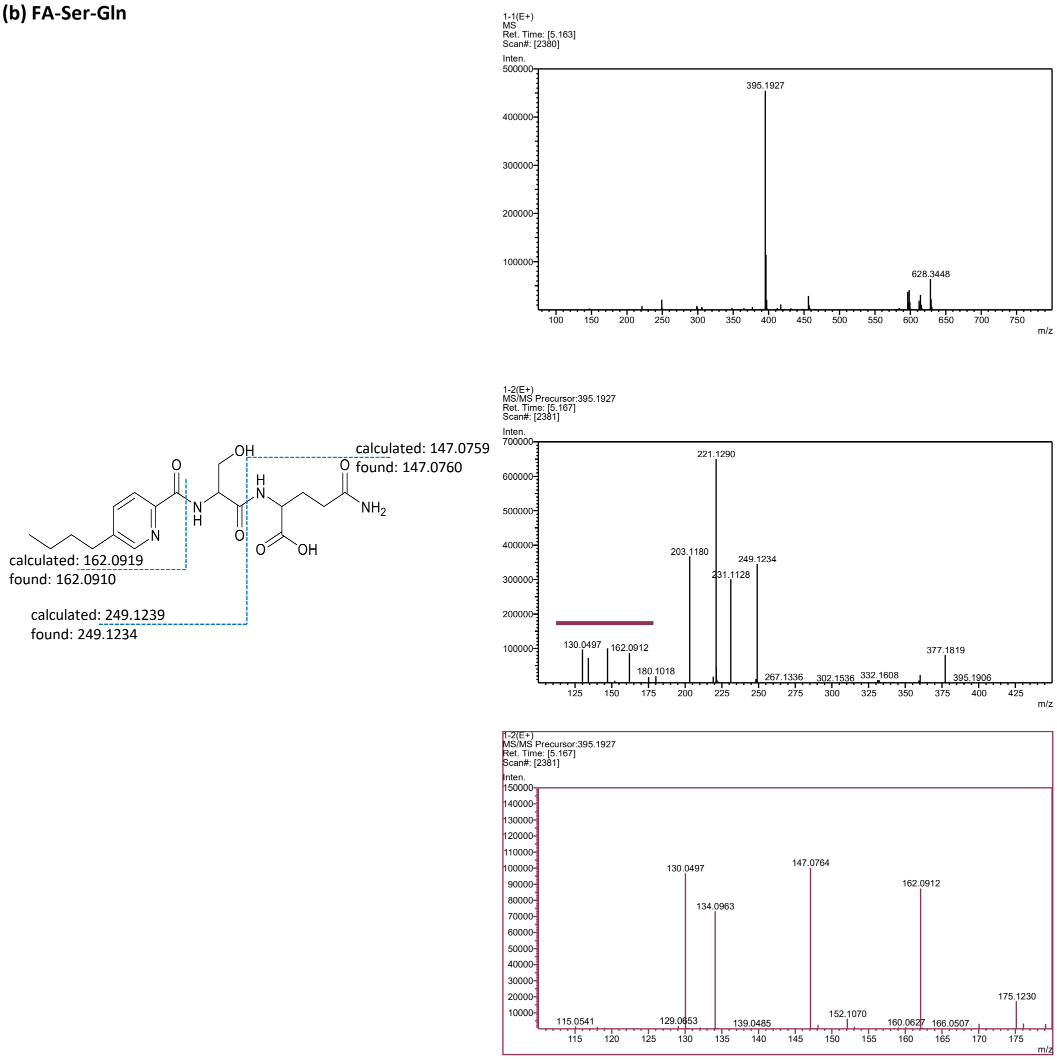

**(c) FA-Ala**

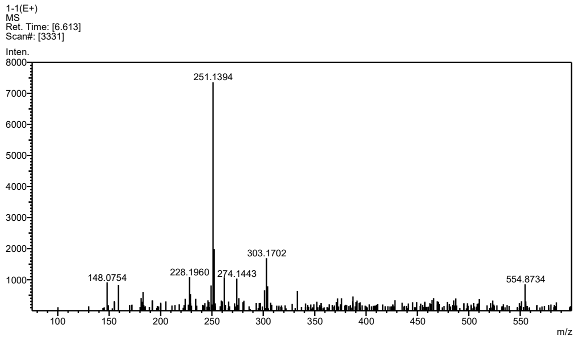


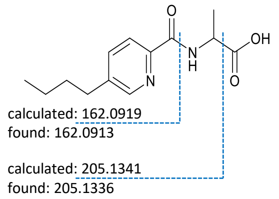


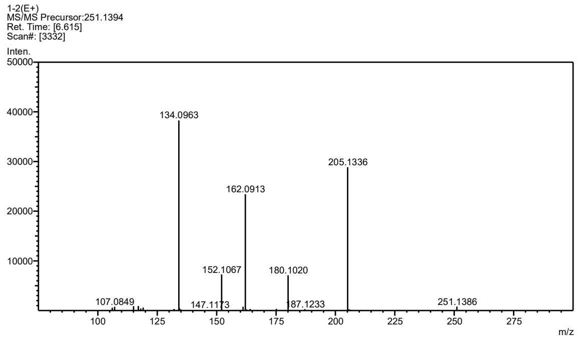


**(d) FA-Ala-Gln**

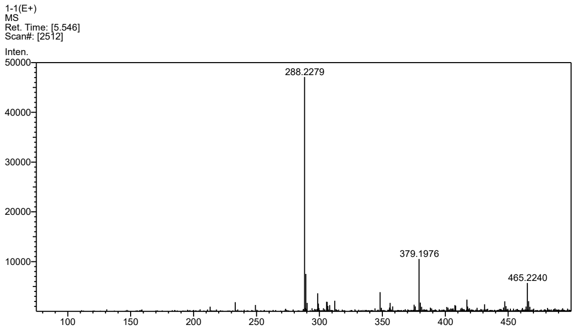


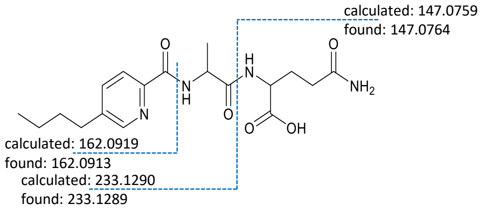


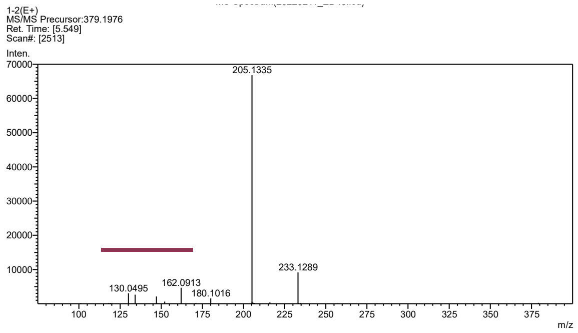


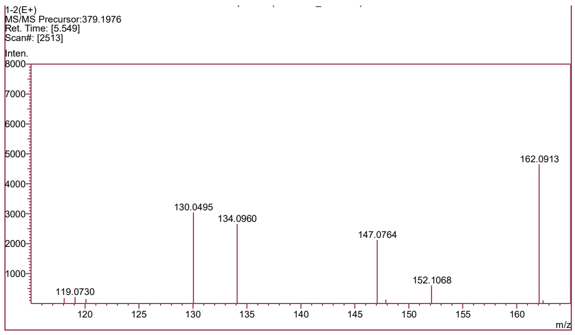

**(e) FA-Thr**

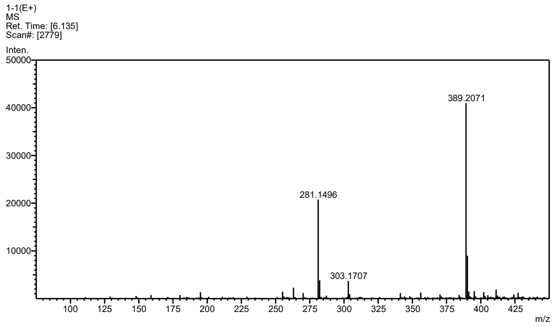


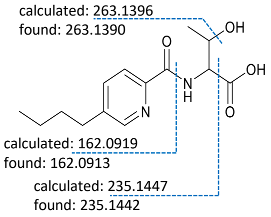


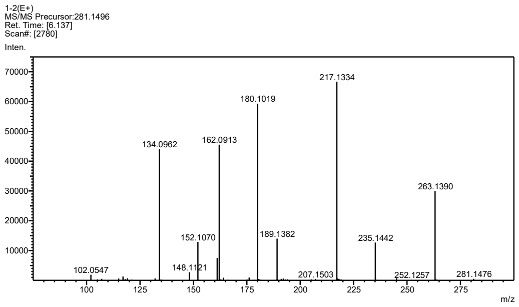


**(f) FA-Gln**

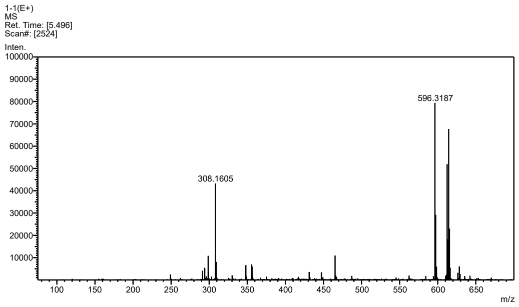


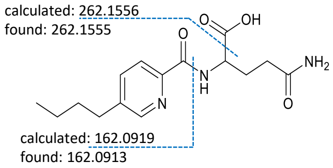


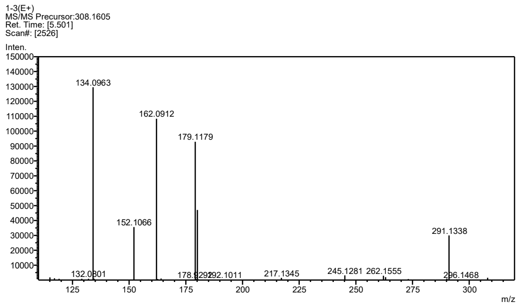

**(g) FA-Gly-Gln**

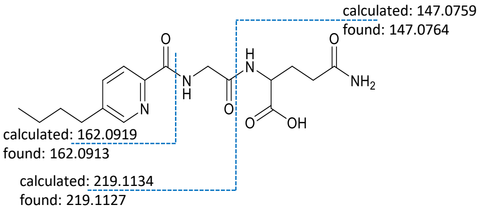


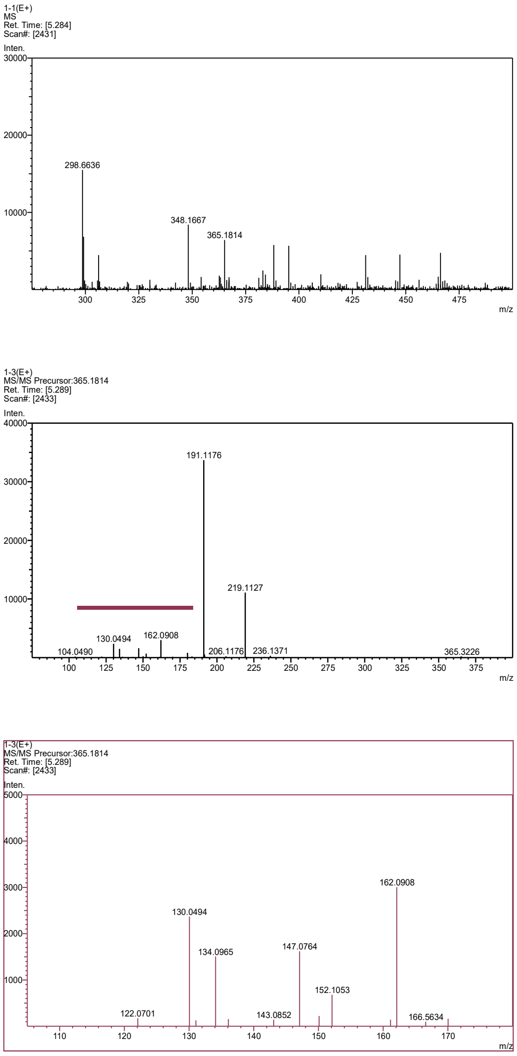


**Fig S6. MS and MS/MS spectra of single and double amino acid conjugates of fusaric acid (FA) detected in *Streptomyces* sp. ATMOS43 culture upon FA treatment**. (+)-HR-ESI-MS spectrum (top panel) and CID MS/MS spectrum of the $[M+H]^+$ ion (one or two bottom panels) of **a)** FA-Serine. **b)** FA-Serine-glutamine. **c)** FA-Alanine. **d)** FA-Alanine-Glutamine. **e)** FA-Threonine. **f)** FA-Glutamine. **g)** FA-Glycine-Glutamine.

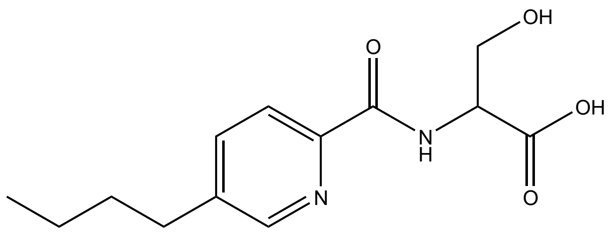


**Fig S7. Chemical structure of FA-Serine synthesized in this work**

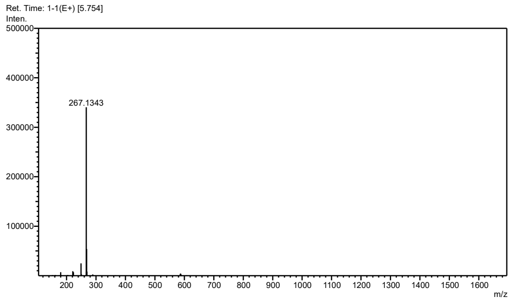


**Fig S8. (+)-HR-ESI-MS spectrum of FA-Serine synthesized in this work**

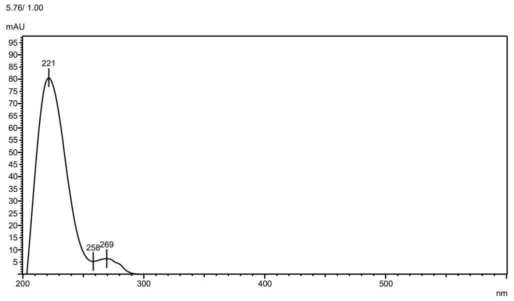


**Fig S9. UV spectrum of FA-Serine synthesized in this work**

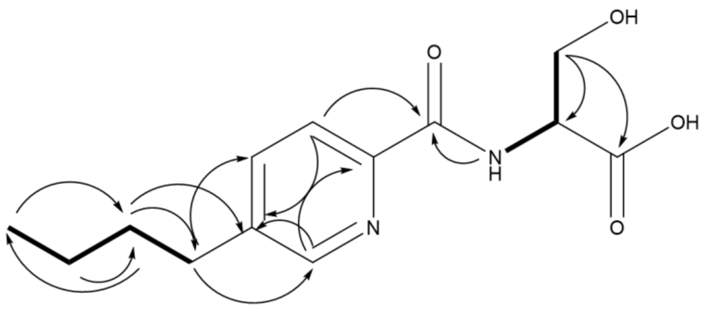


**Fig S10. COSY (▬▬) and HMBC (⟶) correlations of FA-Serine synthesized in this work**

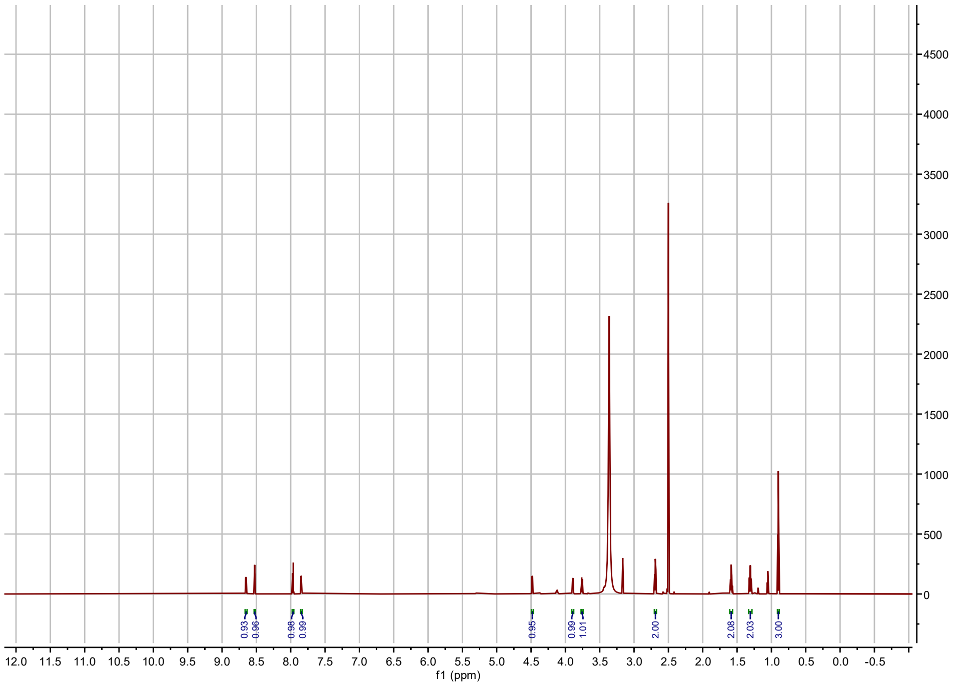

**Fig S11. $^{1}$H-NMR spectrum of Fusaric Acid-Serine synthesized in this work (600 MHz, in DMSO)**

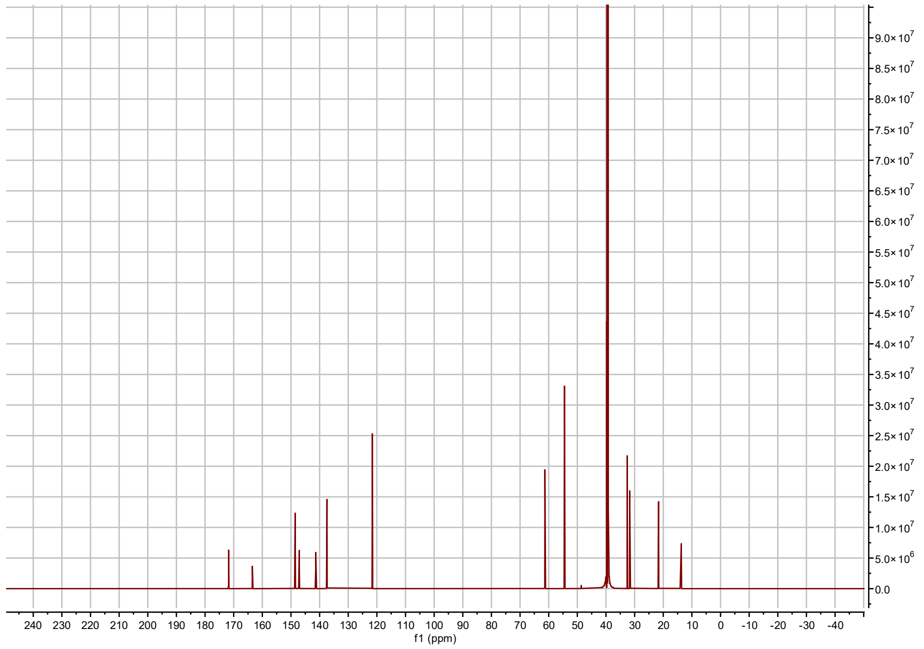

**Fig S12. $^{13}$C-NMR spectrum of Fusaric Acid-Serine synthesized in this work (150 MHz, in DMSO)**

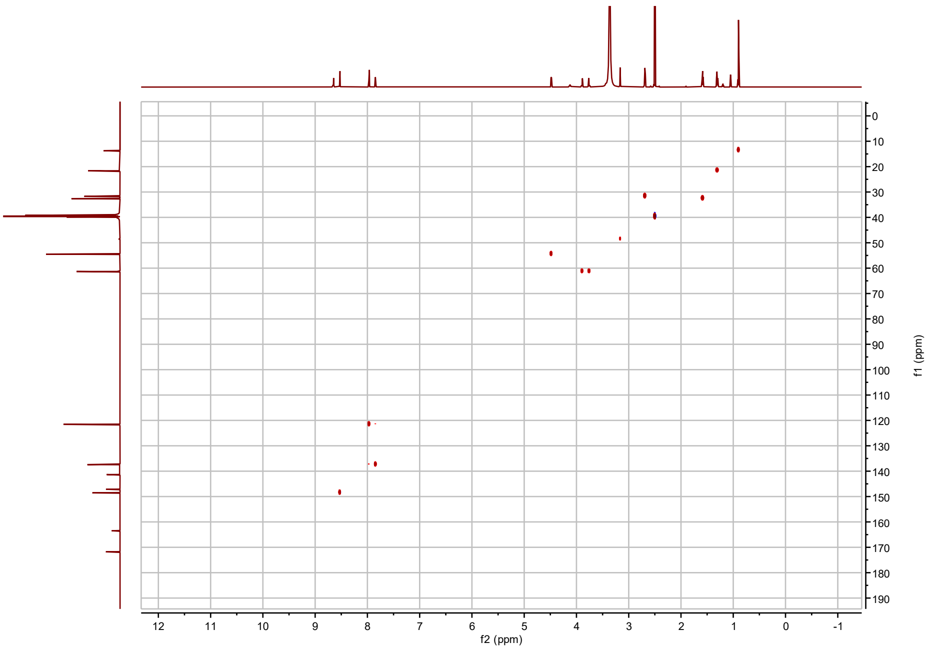


**Fig S13.** $^{1}H$-$^{13}C$ **HSQC spectrum of Fusaric Acid-Serine synthesized in this work (600 MHz, in DMSO)**

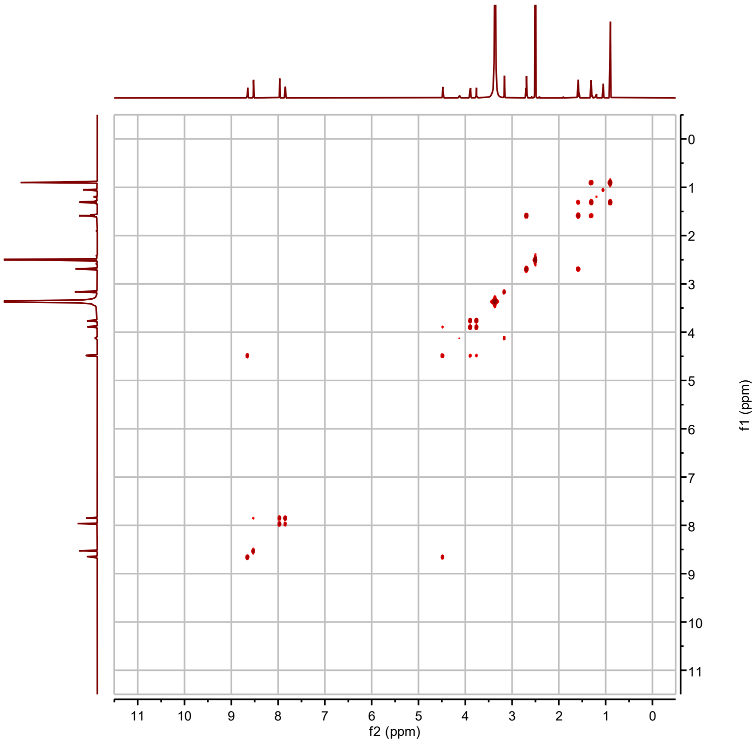


**Fig S14.** $^{1}H$-$^{1}H$ **COSY spectrum of Fusaric Acid-Serine synthesized in this work (600 MHz, in DMSO)**

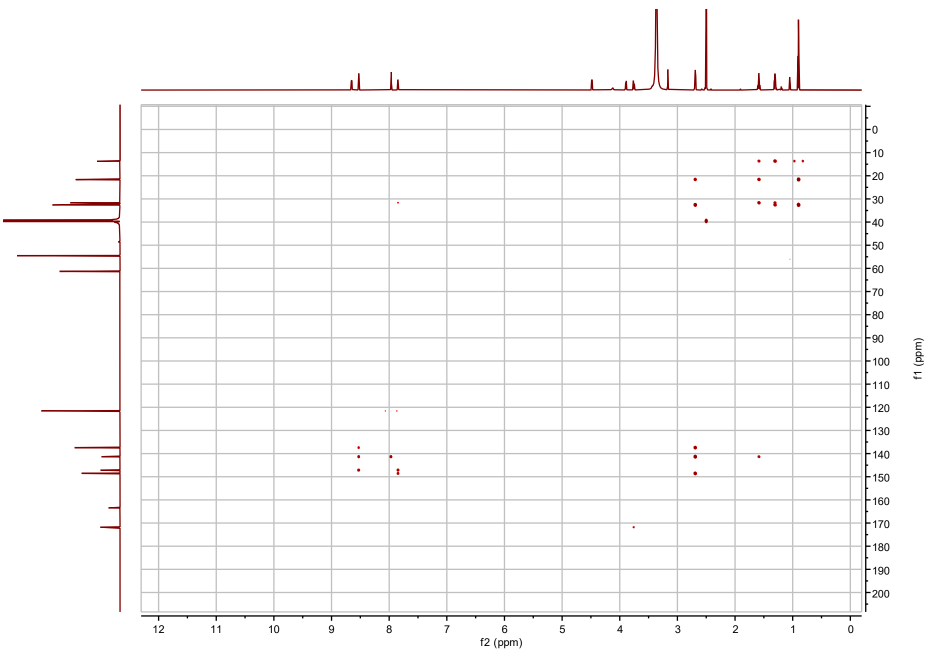


**Fig S15.** $^{1}H$-$^{13}C$ **HMBC spectrum of Fusaric Acid-Serine synthesized in this work (600 MHz, in DMSO)**

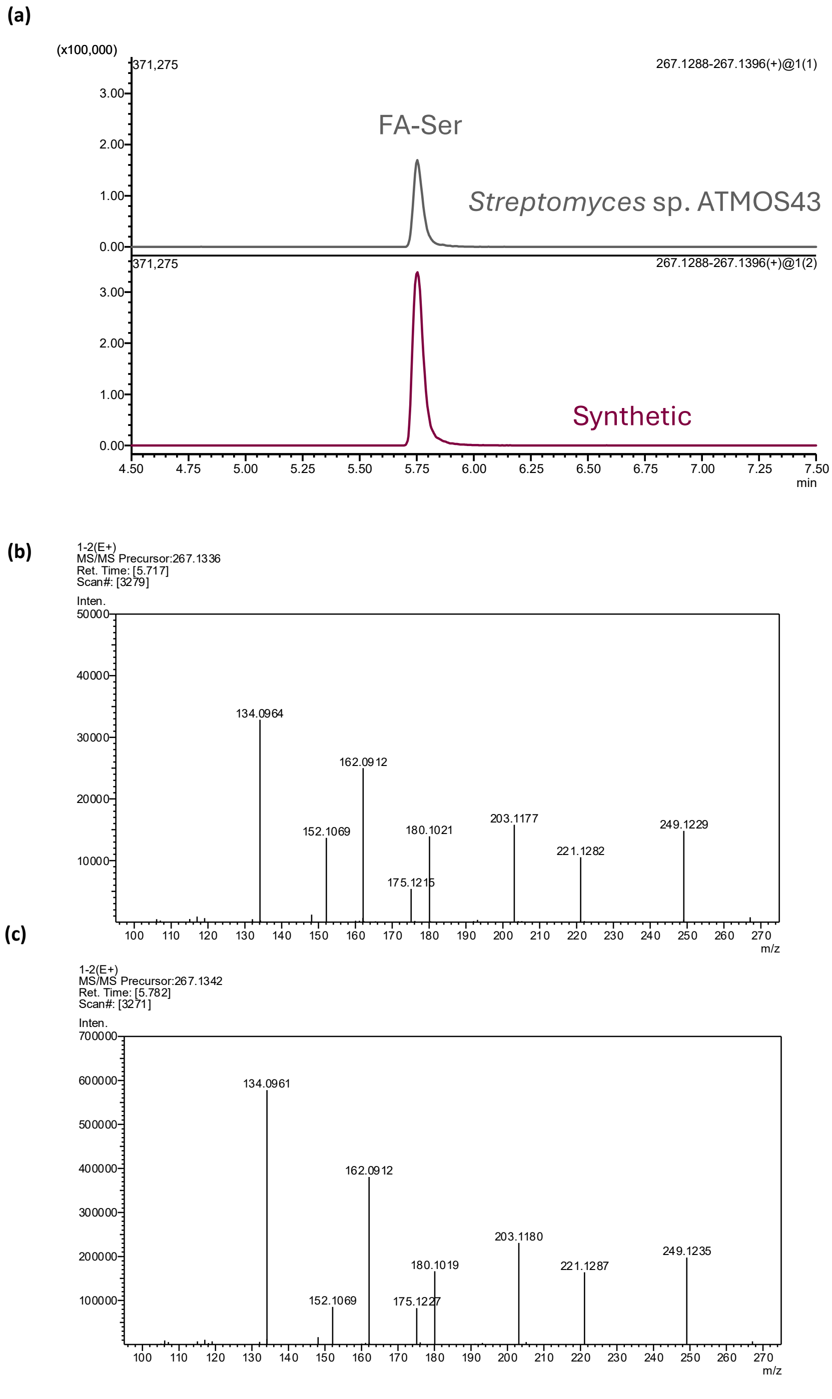


**Fig S16. Confirmation of the identity of the annotated FA-Ser conjugate**. **a)** Extracted Ion Chromatogram (EIC) of FA-Ser from *Streptomyces* sp. ATMOS43 culture extract (top) and synthetic FA-Ser (bottom) **b)** MS/MS spectrum of FA-Ser from *Streptomyces* sp. ATMOS43 culture extract. **c)** MS/MS spectrum of synthetic FA-Ser.

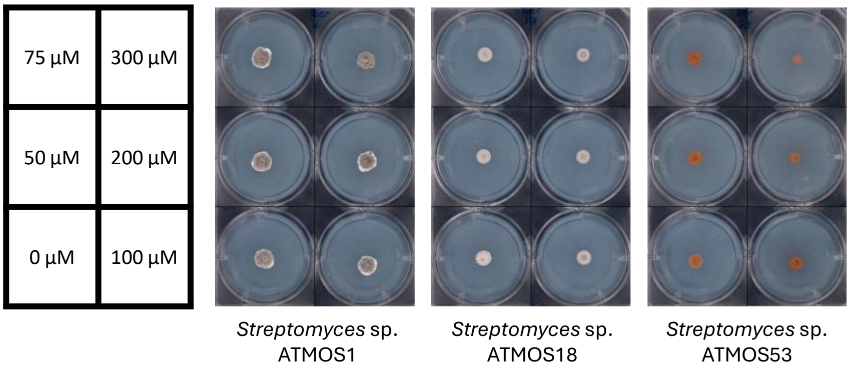


**Fig S17. Minimum Inhibitory Concentration (MIC) assay of FA-Ser across previously tested ATMOS strains**.

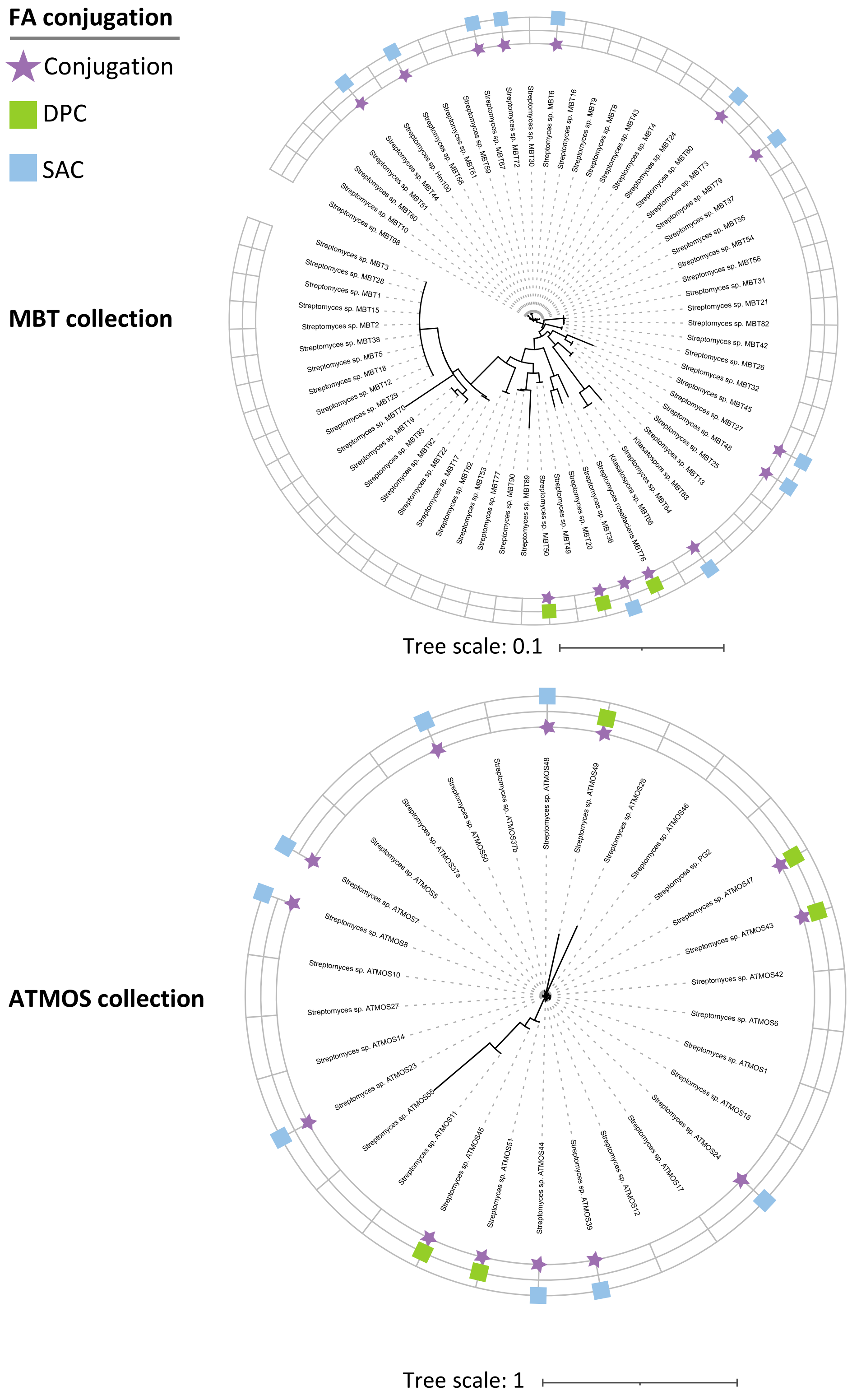


**Fig S18. Phylogenetic tree based on 16S rRNA sequences of the MBT (top) and ATMOS (bottom) actinobacterial strains showing the distribution of single (SAC) and double (DPC) amino acid conjugation activity of fusaric acid.**

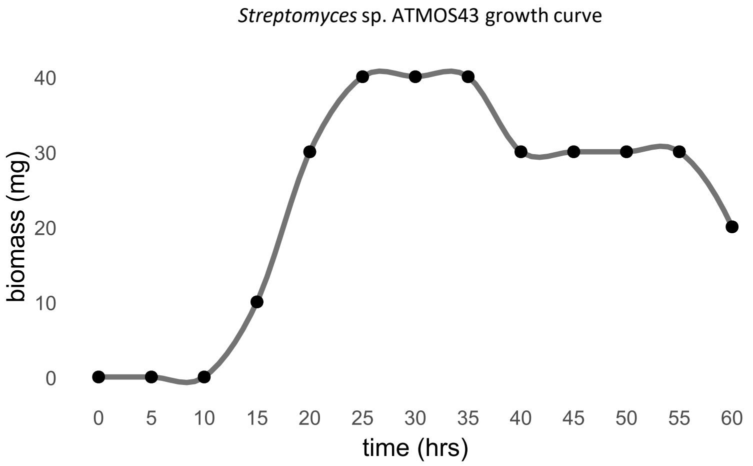


**Fig S19. Growth curve of *Streptomyces* sp. ATMOS43 in ISP2 based on dry biomass.**

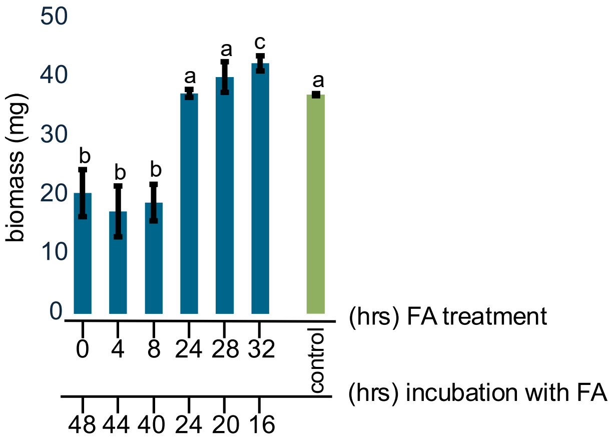


**Fig S20. Biomass accumulation of *Streptomyces* sp. ATMOS43 upon FA feeding at different stages of growth.** FA feeding at early growth stages (0, 4, and 8 h) results in significant growth inhibition compared to control cultures. FA feeding at late growth stages (24, 28, and 32 h) results in similar biomass yields as control cultures. One-way ANOVA, followed by Tukey's HSD test was used to compare growth inhibition levels (n=3). Different letters show statistically significant differences (*p*-value < 0.05).

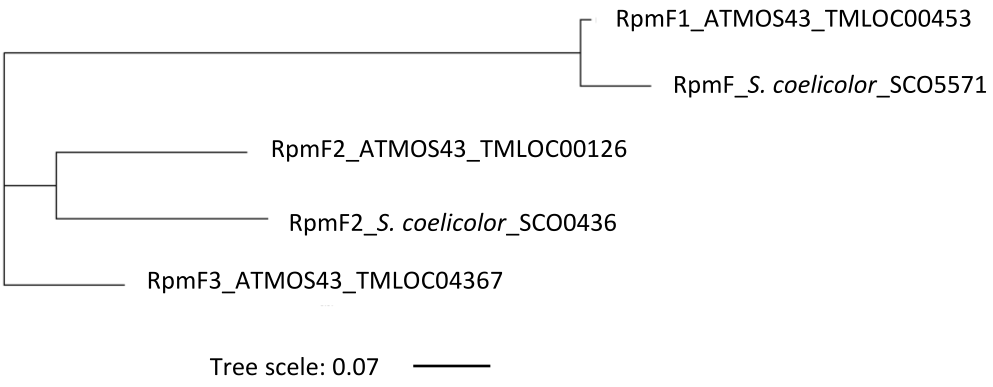


**Fig S21. Phylogenetic tree of L32 ribosomal proteins in *S. coeilicolor* M145 and *Streptomyces* sp. ATMOS43 based on amino acid sequence**

| | |
|---|---|
| 75 µM | 300 µM |
| 50 µM | 200 µM |
| 0 µM | 100 µM |

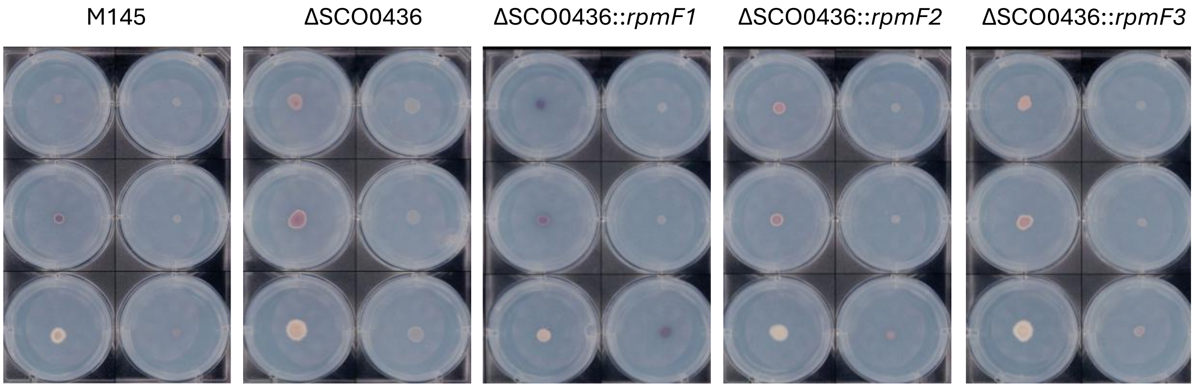


**Fig S22. Minimum Inhibitory Concentration (MIC) assay of FA against *S. coelicolor* M145, its rpmF2 (SCO0436) knock-out mutant, and strains individually complemented with one of the RpmF proteins encoded by *Streptomyces* sp. ATMOS43**. No apparent differential toxicity of FA is observed between the strains. Images show plate bottom.

**(a)**

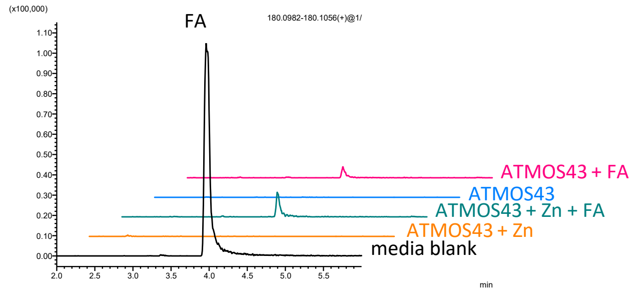


**(b)**

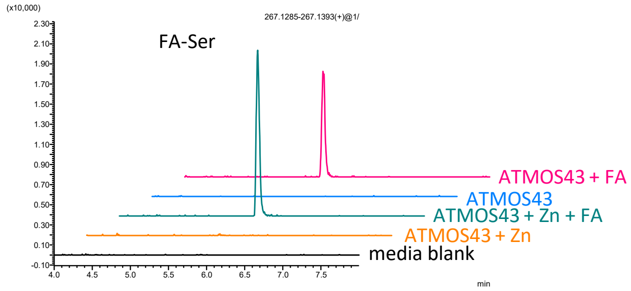


**Fig S23. Zinc supplementation does not inhibit the amino acid conjugation of fusaric acid (FA) by *Streptomyces* sp. ATMOS43. a)** Extracted Ion chromatograms (EIC) showing the depletion of FA in Streptomyces sp. ATMOS43 cultures with and without zinc supplementation. **b)** EIC showing the similar formation of FA-Serine in *Streptomyces* sp. ATMOS43 with and without zinc supplementation.
