## Supplemental Materials and Methods for "Plant-associated *Streptomyces* detoxify the mycotoxin fusaric acid by amino acid conjugation"

**Metabolite profiling using LC-MS/MS**

For liquid chromatography-tandem mass spectrometry (LC-MS/MS) analyses, the dry extracts were dissolved in MeOH to a final concentration of 0.5 mg/mL. LC-MS/MS acquisition was performed using Shimadzu Nexera X2 ultra high-performance liquid chromatography (UPLC) system, with attached photodiode array detector (PDA), coupled to Shimadzu 9030 QTOF mass spectrometer equipped with a standard electrospray ionization (ESI) source unit, in which a calibrant delivery system (CDS) is installed. Samples were injected into a Waters Acquity HSS C18 column (1.8 µm, 100 Å, 2.1 × 100 mm). The column was maintained at 30 °C and run at a flow rate of 0.5 mL/min, using 0.1% formic acid in $H_2O$, and 0.1% formic acid in acetonitrile (ACN) as solvents A and B, respectively. The gradient used was 5% B for 1 min, 5–85% B for 9 min, 85–100% B for 1 min, and 100% B for 4 min. The column was re-equilibrated to 5% B for 3 min before the next run was started. The PDA acquisition was performed in the range of 200–600 nm, at 4.2 Hz, with 1.2 nm slit width. The flow cell was maintained at 40 °C.

All the samples were analysed in positive polarity, using data-dependent acquisition mode. In this regard, full scan MS spectra (m/z 100–1700, scan rate 10 Hz, ID enabled) were followed by two data-dependent MS/MS spectra (m/z 100–1700, scan rate 10 Hz, ID disabled) for the two most intense ions per scan. The ions were fragmented using collision-induced dissociation (CID) with fixed collision energy (CE 20 eV) and excluded for 1 s before being re-selected for fragmentation. The parameters used for the ESI source were: interface voltage 4 kV, interface temperature 300 °C, nebulizing gas flow 3 L/min, and drying gas flow 10 L/min.

**LC-MS based comparative metabolomics and MS/MS-based molecular networking**

The files were imported into Mzmine 2.35 for data processing [1]. Raw data obtained from LC-MS analysis were converted to mzXML centroid files using Shimadzu LabSolutions Postrun Analysis. Mass tolerance was set to 0.002 *m/z* or 15.0 ppm, RT tolerance was set to 0.1 min and noise level was set to 1.2E2. Mass and RT tolerance were set to those parameters for all processing steps unless stated otherwise.

Chromatograms were built using ADAP chromatogram builder (minimum group size in number of scans: 8; group intensity threshold: 1.8E2; minimum highest intensity 5.0E2) [2]. All detected peaks were smoothed (filter width: 9), and the chromatograms were deconvoluted (algorithm: local minimum search; chromatographic threshold: 95%; minimum relative height: 1%; minimum ratio of peak top/edge: 1.8; minimum absolute heigh:1.8E2; peak duration: 0.03-3.00 min). The detected peaks were deisotoped (monotonic shape; maximum charge: 2; representative isotope: most intense). Peak lists of all samples were aligned using join aligner (weight for RT =50; weight for m/z = 50; compare isotopic pattern with a minimum score of 50% and minimum absolute intensity 1.5E2). Improperly detected peaks that were listed in at least one sample in the aligned feature list were filled with the gap filling peak finder algorithm (Intensity tolerance 10 % and RT tolerance: 0.1 min). Subsequently, fragment ions (RT tolerance: 0.05 min, maximum fragment peak height: 50%, and minimum MS2 peak height: 0.0E0), adducts (RT tolerance: 0.05, maximum relative adduct peak height: 5000% for the identification of $[M+Na]^+$, $[M+K]^+$, $[M+NH4]^+$) and complexes (Ionization method: $[M+H]^+$, RT tolerance: 0.05, maximum complex height: 50%) were identified. Duplicate peaks were filtered to remove artifacts caused by detector ringing using old average filter mode (mass tolerance: 1.0 *m/z* or 1000.0 ppm).

The aligned peaks were exported to a GNPS mass-feature quantification table, with the associated MS/MS spectra exported as .mgf files. Further filtering in Excel involved removal of features with consistent peak area detection below 3000 were removed in addition to features originating from culture media. Both the precursor ion and the MS/MS fragment ion mass tolerance were set to 0.02 Da. The minimum cosine score was set to 0.75 and the minimum matched peaks set to 4. MS2LDA analysis was conducting with 200 identified motifs and 5000 iterations. The overlap score of MS/MS spectra to Mass2Motif was set to 0.4. Cytoscape 3.9.1 was used for visualization of the generated molecular network. Metabolomics analysis and statistical analyses at individual feature levels were performed in R, and plots were generated by ggplot2 package in R.

**FA levels in cultures of ATMOS strains**

LC-MS analysis was performed on a Shimadzu LC-20AD/8040 system with a Shimadzu Shim-Pack GISS-HP C18 column (3.0 x 150 mm, 3 µm) at 30 °C and equipped with a UV detector monitoring at 220 and 254 nm. The following solvent system, at a flow rate of 0.5 mL/min, was used: solvent A, 0.1 % formic acid in water; solvent B, acetonitrile. Gradient elution was as follows: 95:5 (A/B) for 1 min, 95:5 to 0:100 (A/B) over 8 min, 0:100 (A/B) for 2 min, then reversion back to 95:5 (A/B) over 1 min, 95:5 (A/B) for 3 min. The injection volume was 10 µL. The ionization source was operated in positive mode, the nebulizing gas set at 3 L/min, the drying gas at 15 L/min, and the desolvation line (DL) temperature at 250 °C. The detection of FA was carried out with multiple reaction monitoring (MRM) of the following transitions: 180 -> 134.

**Proteomics sample preparation and analysis**

Samples were prepared as described in [3, 4] with some modifications. Briefly, a bacterial culture was pelleted and lysed in lysis buffer (4% SDS, 100 mM Tris-HCl pH 7.6, 0.1M DTT, 50 mM EDTA) and disrupted by sonication (Bioruptor Plus, Diagenode). Total protein was precipitated using the chloroform-methanol method [5], and the proteins dissolved in 1% sodium deoxycholate at 95 °C. The protein concentration was measured at this step using BCA method. Protein samples were then reduced by adding 5 mM DTT and incubating at 60 °C for 30 min, followed by thiol group protection with 21.6 mM iodoacetamide incubation at room temperature in dark for 30 min. Then 0.1 µg trypsin (recombinant, proteomics grade, Roche) per 10 µg protein was added, and samples were digested at 37 °C for 8 h and then at room temperature. After digestion, trifluoroacetic acid was added to 1% and samples were incubated at 37 °C for 30 min followed by centrifugation to precipitate deoxycholate. Peptide solution containing 6 µg peptide was then cleaned and desalted using STAGE-Tips [6]. Briefly, 6 µg of peptide was loaded on a conditioned StageTip with C18 disk (AttractSPE Tips C18.T1.10.960, Affinisep), washed once with 0.5% formic acid solution, and eluted with 80% acetonitrile and 0.5% formic acid. Acetonitrile was then evaporated in a SpeedVac. Final peptide concentration was adjusted to 40 ng/µL using sample solution (3% acetonitrile, 0.5% formic acid) for analysis.

Peptides were separated via nanoflow reversed-phase liquid chromatography using a nanoElute 2 LC system (Bruker Daltonics) coupled to a timsTOF HT mass spectrometer (Bruker) with 0.1% FA (solution A) and 0.1% FA/99.9% ACN (solution B) as the mobile phases. The samples were loaded on a trap column (PepMap $C_{18}$, 5 mm x 0.3 mm, 5 µm, 100 Å, Thermo Scientific) followed by elution and separation on the analytical column (PepSep $C_{18}$, 25 cm x 75 µm, 1.5 µm, 100 Å, Bruker) kept at 50 °C using a gradient of 3 - 25% solvent B in 25 min, 25 - 32% B in 5 min, 32 - 95% B in 5 min and 95% B for 10 min at a flow rate of 300 nL/min. Peptides were introduced to a TimsTOF HT (Bruker Daltonics) using a 20 µm ID fused silica emitter (Bruker Daltonics) installed in a nano-electrospray ion source (CaptiveSpray source, Bruker Daltonics) with spray voltage set to 1400 V.

For DIA acquisition, peptides were analysed with a TimsTOF HT (Bruker Daltonics) running in DIA-PASEF mode. The DIA-PASEF method was optimized for the specific sample type using the py_diAID tool [7]. The method covered an ion mobility range from 1.35 to 0.7 Vs cm-2 and an *m/z* range of 300 to 1300, using 10 DIA-PASEF scans with two isolation windows per scan, resulting in a cycle time of 1.1 s. Collision energy was linearly decreased from 59 eV at 1.6 Vs cm-2 to 20 eV at 0.6 Vs cm-2. For all experiments the ion mobility dimension was calibrated linearly using three selected ions of the Agilent ESI LC-MS Tuning Mix [m/z, 1/K0: (322.0481, 0.7318 Vs cm−2), (622.0289, 0.9848 Vs cm−2), (922.0097, 1.1895 Vs cm−2)]. Mass calibration was performed with sodium formate in HPC mode.

DIANN (Academia v 2.1.0) [8, 9] is used in raw data processing. Spectrum library is generated in-silico from the proteome generated from genome sequence, with parameters: 1% FDR cutoff; mass accuracy 15 ppm; precursor *m/z* range 280–1800; fragment *m/z* range 200–1800; peptide length 7–30; precursor charge 1–4; loose trypsin digestion (cut all after K and R); allow 1 missed cleavage; fixed modification is set to C carbamidomethylation; allowing 1 variable modification methionine oxidation or N-term methionine excision; peptidoform confidence scoring enabled. Raw data was then searched against the generated spectrum library with match between runs enabled.
